# Liquid droplet aging and fibril formation of the stress granule protein TIA1 low complexity domain

**DOI:** 10.1101/2022.03.30.486471

**Authors:** Yuuki Wittmer, Khaled M. Jami, Rachelle K. Stowell, Truc Le, Ivan Hung, Dylan T. Murray

**Affiliations:** Department of Chemistry, University of California, Davis, California 95616, USA; National High Magnetic Field Laboratory, Tallahassee, Florida 32310, USA

**Keywords:** TIA1, T-cell restricted intracellular antigen-1, solid state nuclear magnetic resonance, biomolecular condensation, liquid-liquid phase separation, liquid droplet, low complexity domain, amyotrophic lateral sclerosis, frontotemporal dementia, protein fibril, gelation

## Abstract

Protein domains biased toward a few amino acid types are vital for the formation of biomolecular condensates in living cells. These membraneless compartments are formed by molecules exhibiting a range of molecular motions and structural order. Missense mutations increase condensate persistence lifetimes or structural order, properties that are thought to underlie pathological protein aggregation. We examined seeded fibrils of the T-cell restricted intracellular antigen-1 low complexity domain and determined residues 338–357 compose the rigid fibril core. Aging of wild-type and P362L mutant low complexity domain liquid droplets resulted in fibril assemblies that are structurally distinct from the seeded fibril preparation. The results show that most disease mutations lie outside the region that forms homogeneous fibril structure, the droplets age into conformationally heterogenous fibrils, and the P362L disease mutation does not favor a specific fibril conformation.

## INTRODUCTION

Granular condensates of proteins and nucleic acids are integral to RNA metabolism.These membraneless organelles facilitate RNA processing and transport, and control proteostasis during cellular stress.^1,2^ Macroscopic, fluorescence based measurements on cultured cells show that these RNA granules contain a heterogenous collection of molecules undergoing varying levels of molecular motion.^3^ Experiments using purified proteins reveal that the condensation of homogeneous solutions of specific proteins into liquid droplet structures reproduce the macroscopic behavior of *in vivo* RNA granules.^4,5^ A variety of weak, multivalent interactions facilitate this behavior, which is dictated by the properties of the specific proteins involved.^6^

A low complexity (LC) domain is a protein segment that is biased towards a subset of the 20 amino acid types typically used for natural protein synthesis.^7^ LC domains are well-represented in proteins that form a variety of macroscopic assemblies in living cells^8^ and are of significant interest in the context of biomolecular condensation. These sequences inherently contain the multivalent property required for liquid droplet formation^9^ and can assemble into more rigid amyloid-like fibrils functionally and pathologically.^10,11^ The gelation, or aging, of liquid droplets, which are normally characterized by significant molecular motion and disorder, is particularly pertinent for understanding how RNA granules fail to disassemble, facilitating the buildup of protein aggregates.^4^

The misassembly of LC domain proteins is intricately linked to neurodegenerative disease. Amyotrophic lateral sclerosis (ALS) is a progressive neurodegenerative disease of the spinal cord and brain that results in the loss of muscle movement.^12^ Frontal temporal dementia (FTD) is the second most common form of dementia after Alzheimer’s disease.^13^ These two diseases have many genetic links to LC domain proteins involved in the formation of RNA granules.^14^ The RNA-binding protein T-cell-restricted intracellular antigen-1 (TIA1), has ALS and FTD missense mutations clustered in a C-terminal LC domain.^15^ While the buildup of TIA1-rich inclusions is not typical of ALS and FTD pathology, TIA1 disease mutations result in the persistence and gelation of stress-related RNA granules, and are associated with the pathological deposition of other LC domain proteins.^15^

Functional activity of the TIA1 protein includes the regulation of RNA splicing and translation.^16^ The TIA1 LC domain is a 96 amino acid sequence biased toward Gln, Gly, Tyr, and Pro residues that assembles into amyloid-like fibrils and liquid droplets in vitro.^15,17–19^ Organized and reversible fibrillar self-assembly is possibly a functional activity for the domain.^20^ Yet, despite significant interest in TIA1 mediated condensation processes, there are few structural characterizations of TIA1 fibril formation. The P362L LC domain mutation delays the disassembly of functional full length TIA1 assemblies.^17^ Several Pro-to-hydrophobic mutations in the TIA1 LC domain lead to increased aggregation rates for the protein and suggest an antipathogenic role for Pro residues.^17^ However, the residue-specific effects of LC domain mutations on the gelation of TIA1 liquid droplets remain uncharacterized at high resolution. The biological function and pathogenicity of TIA1 involves LC domain self-assembly processes for which the molecular mechanisms remain unknown.^20^

Here we report the results from solid state nuclear magnetic resonance (NMR) measurements, electron and bright field microscopy imaging, and fluorescence assays that characterize condensed states of wild-type and P362L mutant forms of the TIA1 LC domain. High-resolution measurements of seeded fibrils report on the precise location of the most fibril-prone region of the protein. Analysis of aged liquid droplets provide insight into the structural changes underlying liquid droplet gelation and how an ALS and FTD mutation affects the process. Our results provide a basis for understanding the sequence context of disease mutations in the TIA1 LC domain as it relates liquid droplet aging and a foundation for understanding the broader conformational space sampled by the TIA1 LC domain during proper biological function and disease pathology.

## RESULTS

### TIA1 LC Domain Fibril Seeding

Figure 1A shows the domain structure of the wild-type full-length TIA1 protein. Three N-terminal RNA-binding motifs (RRM, RNA-recognition motif) are followed by a 96-residue LC domain. The primary sequence of the TIA1 LC domain shown in Figure 1B reveals the LC domain is biased toward a subset of the 20 most common amino acids and is dominated by 22% Gln, 16% Gly, 12% Pro, 9% Tyr, and 9% Asn. These residues are well-distributed across the LC domain. LC domain mutations associated with neurodegenerative disease are colored red in Figure 1B.^15^ These mutations are dispersed along the length of the TIA1 LC domain rather than localized to a specific region of the primary sequence. Ala, Ile, Leu, Met, Val, Trp, Ser, Thr residues are less common in the TIA1 LC domain. Of these, Ala and Val residues are found throughout the LC domain and Trp is spread throughout the first two-thirds of the LC domain, while the Ile residues are confined to the N-terminal region and Ser and Thr residues are located at the C-terminal region. The distribution of the less common amino acids makes them highly informative reporters on structure and dynamics in the LC domain.

**Figure 1:**
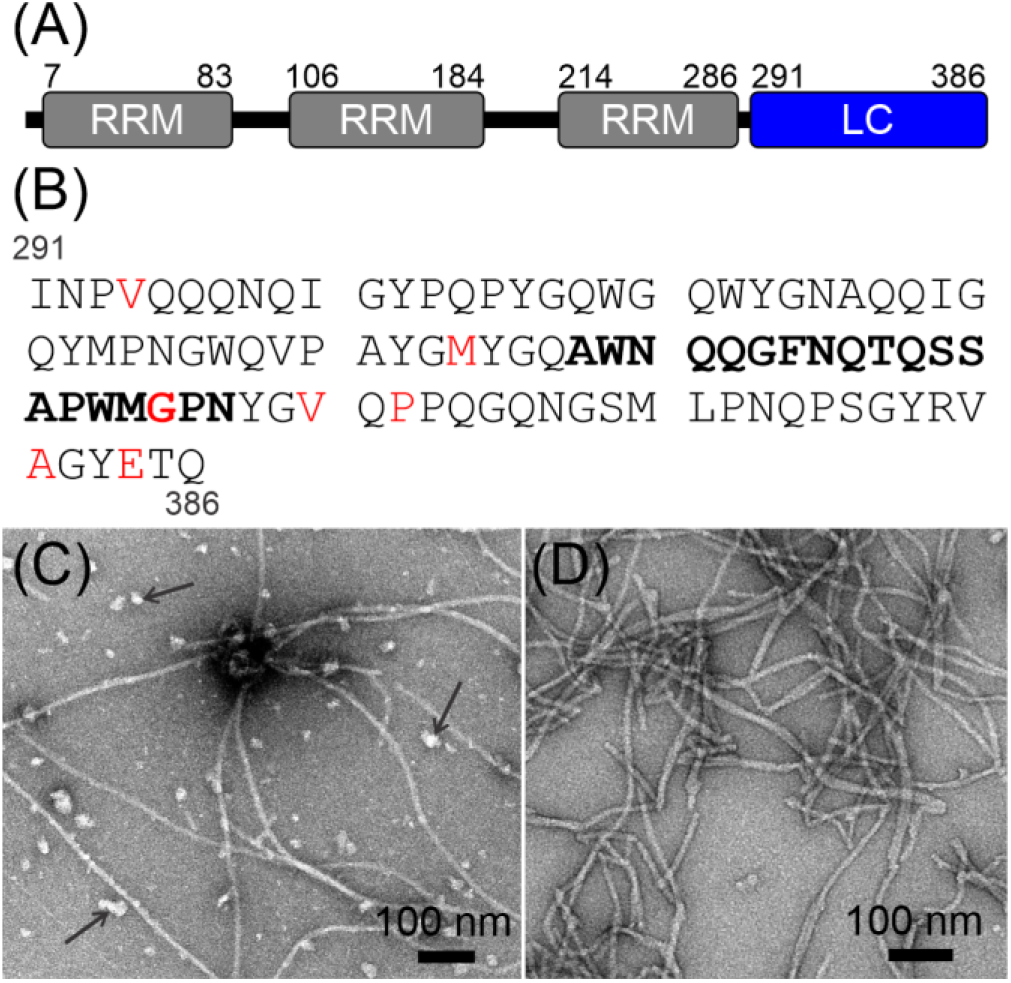
TIA1 Domain Structure and Fibril Seeding. (A) The TIA1 protein is composed of three RNA-recognition motifs (RRM) that precede a 96-residue LC domain. (B) The TIA1 LC domain primary sequence in single letter amino acid code. The locations of missense mutations linked to neurodegenerative disease are highlighted in red. The seeded fibril core forming segment determined in this work is in bold letters. (C) A negatively stained TEM image of TIA1 LC domain fibrils formed quiescently without any seeding. Arrows point to amorphous aggregates that form concomitantly with the fibrils. (D) A negatively stained TEM image of the seeded TIA1 LC domain fibrils shows a reduced number of amorphous aggregates.

A seeding procedure similar to that used for the β-amyloid peptide^21^ was used to obtain fibrils of TIA1 LC domain. The negatively stained transmission electron microscopy (TEM) image in Figure 1C shows that after extensive removal of denaturant and quiescent incubation, the TIA1 LC domain forms both amorphous and fibrillar aggregates in a low ionic strength and near neutral pH solution. Figure 1D shows that a seeding process in the same buffer conditions reduces the prevalence of the amorphous aggregates, producing solutions of TIA1 LC domain that contain long, thin, and somewhat bundled fibrils. These TIA1 LC domain fibrils appear similar to those from a previously published report,^17^ although differences in staining and image quality prevent any further comparison. The residual soluble protein concentration for this seeded preparation was 1.4 μM. Using a model where fibrils do not fragment,^22^ this concentration corresponds to a Gibbs energy of dissociation (ΔG) for a TIA1 LC domain monomer from the fibril of ∼32 kJ/mol at ∼283 K.

### Residues A338–N357 of the TIA1 LC Domain Form β-strand Rich Protein Fibrils

The cross polarization-based solid state NMR spectra of the seeded TIA1 LC domain fibrils in Figure 2 exhibit a mixture of sharp, resolved signals and regions of either highly overlapped or broad signal intensity. Signals in these spectra arise from immobilized amino acids in the fibril structure. The 2D carbon-carbon correlation spectrum^23,24^ in Figure 2A contains sharp aliphatic signals for Thr, Ser, Ala, Asn, and Asp residues. The spectral regions corresponding to Gln, Glu, Phe, Tyr, Trp, His, Met, Arg, and Leu residues contain broad intensities which arise from either conformational heterogeneity of single residues, molecular motions, or the presence of multiple well-ordered residues. There are also broad resonances for all sidechain carbons of Pro residues. The aromatic region of this spectrum does not exhibit strong or sharp signals and is missing intensity that can be associated with His sidechains. The spectrum also contains signal intensities consistent with Val and Ile residues. Notably, the signals corresponding to CA-CB correlations for the Val and Ile residues are significantly weaker than the signals corresponding to CB-CG and other terminal sidechain correlations, which suggest that the backbone atoms for the Val and Ile residues are not as well ordered as the sidechains.

**Figure 2:**
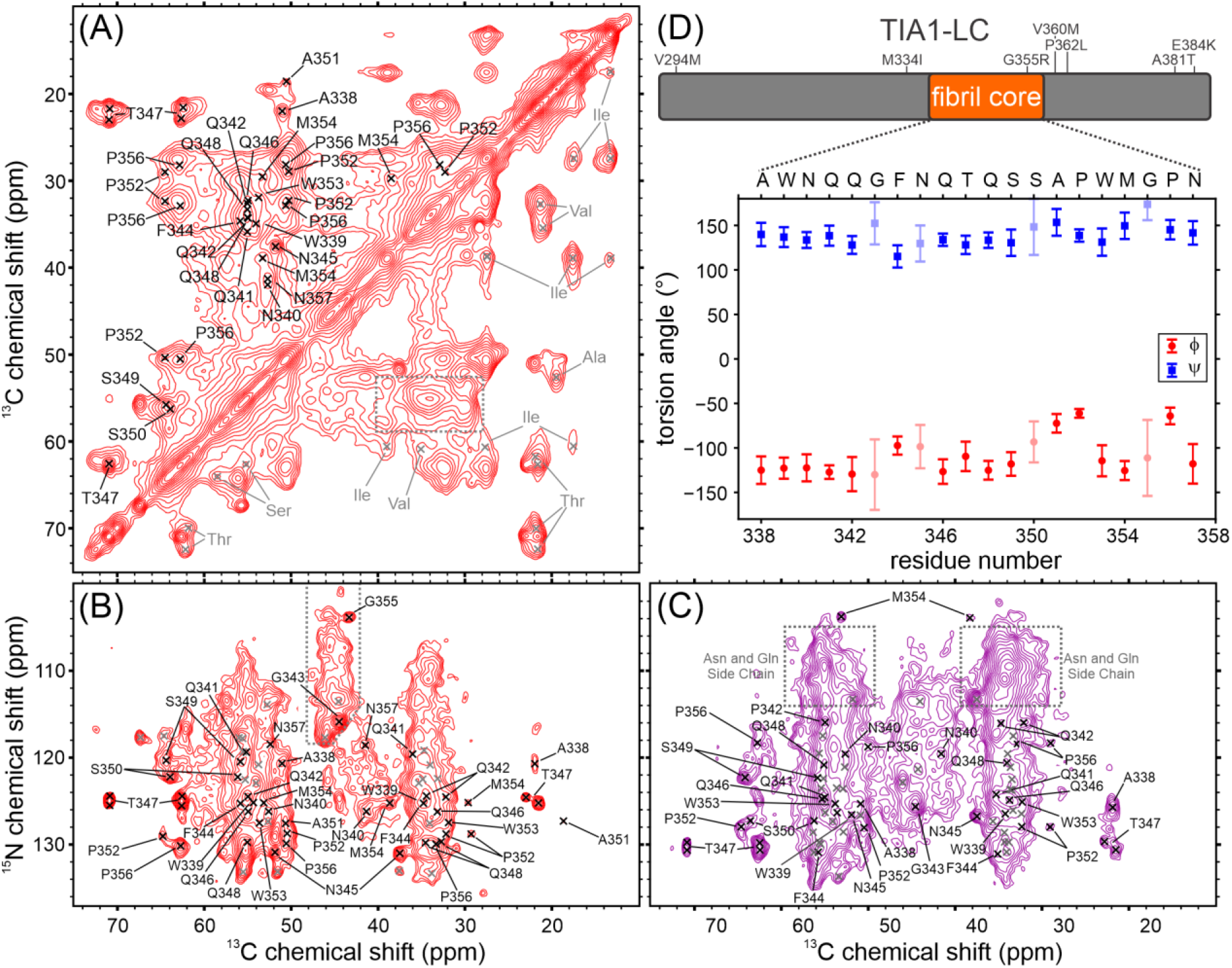
Solid state NMR Characterization of Seeded TIA1 LC Domain Fibrils. (A) The aliphatic region of a carbon-carbon cross polarization-based correlation spectrum of seeded TIA1 LC domain fibrils. The spectrum is pseudosymmetric about the diagonal. Residues labeled in black in the top left of the spectrum were sequence-specifically assigned in this work. Residues labeled in gray in the bottom right of the spectrum were not observed in nitrogen-carbon cross-polarization-based spectra. The gray dashed box highlights the overlapped signal intensity for Gln, Glu, Phe, Tyr, Trp, His, Met, Arg, and Leu residues. (B) The aliphatic region of a nitrogen-carbon cross polarization-based spectrum showing intraresidue correlations. Residues labeled in black were unambiguously assigned in this work. Residues marked in gray were not unambiguously assigned in this work. The gray dashed outline highlights broad signal intensity arising from Gly residues. (C) The aliphatic region of a nitrogen-carbon cross polarization-based correlation spectrum showing interresidue correlations. Residues labeled in black were unambiguously assigned in this work. Residues marked in gray were not unambiguously assigned in this work. The signal intensities indicated with gray dashed boxes originate from nitrogen atoms in Gln and Asn sidechains. (D) A map of the TIA1 LC domain with the locations of the rigid fibril core and disease mutations, and a plot showing the backbone torsion angles predicted from the assigned NMR chemical shifts. Error bars in the plot represent the standard deviation of the predictions and lighter symbols indicate predictions with high uncertainty.

The 2D cross polarization-based nitrogen-carbon spectrum^25^ in Figure 2B reports on rigid Gly residues in the TIA1 LC domain. Two sharp signals are consistent with at least two well-ordered Gly residues. Additional weak and broad signal intensity in this area is consistent with Gly residues that are either undergoing molecular motion or exist in a spectrum of heterogeneous conformations. Much of the remainder of the spectrum is highly overlapped, but sharp signals can be identified for two Thr residues, three Ala, four Ser, and three Asn or Asp residues due to their unique NMR CA and CB chemical shift ranges. The narrow linewidths of these signals indicate they are rigid with well-defined molecular conformations. Pro residues lack an amide proton and therefore do not give rise to strong signals in the spectrum in Figure 2B. The TEDOR experiment^26^ ensures accurate reporting of all rigid Pro residues in nitrogen-carbon spectra, as it uses magnetization transfers originating from the CA proton rather than the amide proton. The TEDOR spectrum of the TIA1 LC domain fibrils in Supplemental Figure S1A contains two sharp and distinct Pro signals that arise from sites that are in well-defined and rigid conformations. There is additional broad signal intensity consistent with Pro residues that are either loosely ordered or sample heterogeneous conformations. The remainder of the TEDOR spectrum is otherwise consistent with the spectrum in Figure 2B.

The proton-carbon spectrum shown in Supplemental Figure S1B uses scalar magnetization transfers^27^ that arise from highly mobile regions of the fibril structure. There is a signal in the spectrum with random coil NMR chemical shifts that can be unambiguously assigned to a Thr CB site. Signals from aromatic atoms in His sidechains are also present in the spectrum. The remainder of the signals in the spectrum arise from other aliphatic carbons that cannot be unambiguously assigned.

A comparison of the amino acid types observed in the spectra in Figures 2A, 2B, and Supplemental Figure 1A with the protein sequence in Figure 1B reveals that the Ala, Gln, Pro, Trp, and Met signals arising from immobilized residues in the fibril structure are potentially spread along the entire TIA1 LC domain. However, the increased mobility of the backbone atoms for Ile and Val residues (Figure 2A) and their location in the N-terminal region of the TIA1 LC domain suggest that this region is not as uniformly ordered or immobilized as the C-terminal region, which is populated with Ala, Ser, and Thr residues that give rise to relatively sharp signals arising from well-ordered and rigid sites in the TIA1 LC domain. The small number of signals in the scalar-based spectrum in Supplemental Figure S1B indicate that very few sites in the TIA1 LC domain are highly mobile. In our TIA1 LC construct, the only His residues and two additional Thr residues are present in the His-tag (see Methods) used to express and isolate the TIA1 LC domain. The lack of strong signals from His residues in the cross polarization-based spectra indicate that these sites are not part of the rigid fibril core. Furthermore, the unambiguous signals from His sidechain sites and a Thr CB site in the spectrum in Supplemental Figure S1B reinforce the interpretation that the tag does not influence the structure of the rigid fibril for the TIA1 LC domain. The 2D spectra presented in Figure 2A, 2B, S1A, and S1B are therefore consistent with a TIA1 LC domain fibril structure formed by a strongly immobilized and well-ordered core composed of but not limited to Ala, Thr, Ser, and Asn or Asp residues, with the remainder of the TIA1 LC domain exhibiting structural heterogeneity and limited motion, and the His-tag undergoing more rapid motion in a random coil-like configuration.

A total of 33 residues were resolved using 3D cross-polarization based spectra. The signal to noise and resolution in these spectra are shown using representative 2D planes in Supplemental Figures S3–5. Almost all spectral regions exhibit well-resolved signal intensities in the 3D spectra, although there is some overlap in the region reporting on CB and CG sites from Gln, Glu, Phe, Tyr, Trp, His, Met, Arg, and Leu residues. However, there are enough features to identify signals from 15 distinct residues. Additionally, signals unambiguously attributable to two Ala, three Gly, two Pro, five Asn or Asp, two Thr, and four Ser residues are clearly identified in these spectra. Signals for Leu, Val, or Ile are not observed in the 3D spectra. All signals identified in these 3D spectra are listed in Supplemental Tables S2-4. These 3D spectra include an NCOCX experiment that provides the interresidue correlations required to associate the observed signals with specific residues in the TIA1 LC domain sequence. The spectrum from a 2D version of this experiment is shown in Figure 2C.

Unambiguous and statistically significant sequence-specific assignments for residues 338–357 were obtained using the MCASSIGN Monte-Carlo simulated annealing algorithm.^28^ The procedure used is similar to our work on other LC domain proteins.^29–32^ A complete description of the assignment calculations is presented in the Methods section. Signals representing an amino acid sequence of XGXX (consistent with residues W309–W312 or I319– Y322) could not be unambiguously assigned due to the low signal to noise of the flanking residues, suggesting an ordered region surrounded by more mobile segments of the protein. Figure 2D shows the TALOS-N^33^ torsion angle predictions from the assigned carbon and nitrogen chemical shifts. The large positive φ and large negative ψ values indicate the immobilized residues are in β-strand conformations. The spectra in Figure 2 and Supplemental Figures S1–S4 therefore are consistent with a β-strand rich fibril structure formed by a 20-residue core composed of residues 338–357 flanked regions that are not highly mobile and lack well-defined structure.

### Both Wild-type and P362L Mutant TIA1 LC Domain Liquid Droplets Age into Protein Fibrils

The bright field microscope images in Figure 3A show that removal of denaturant from purified solutions of wild-type and P362L mutant TIA1 LC domain at near neutral pH and moderate ionic strength produces solutions containing liquid droplets. The droplets occur when the urea concentration drops below 1.5 M during dialysis^32^ and persist for up to 4 h. Supplemental movie S1 shows that at 1.5 h, when the denaturant concentration is ∼0.8 M, the droplets fuse with one another, consistent with a condensed liquid droplet phase of mobile TIA1 LC domain protein.

**Figure 3:**
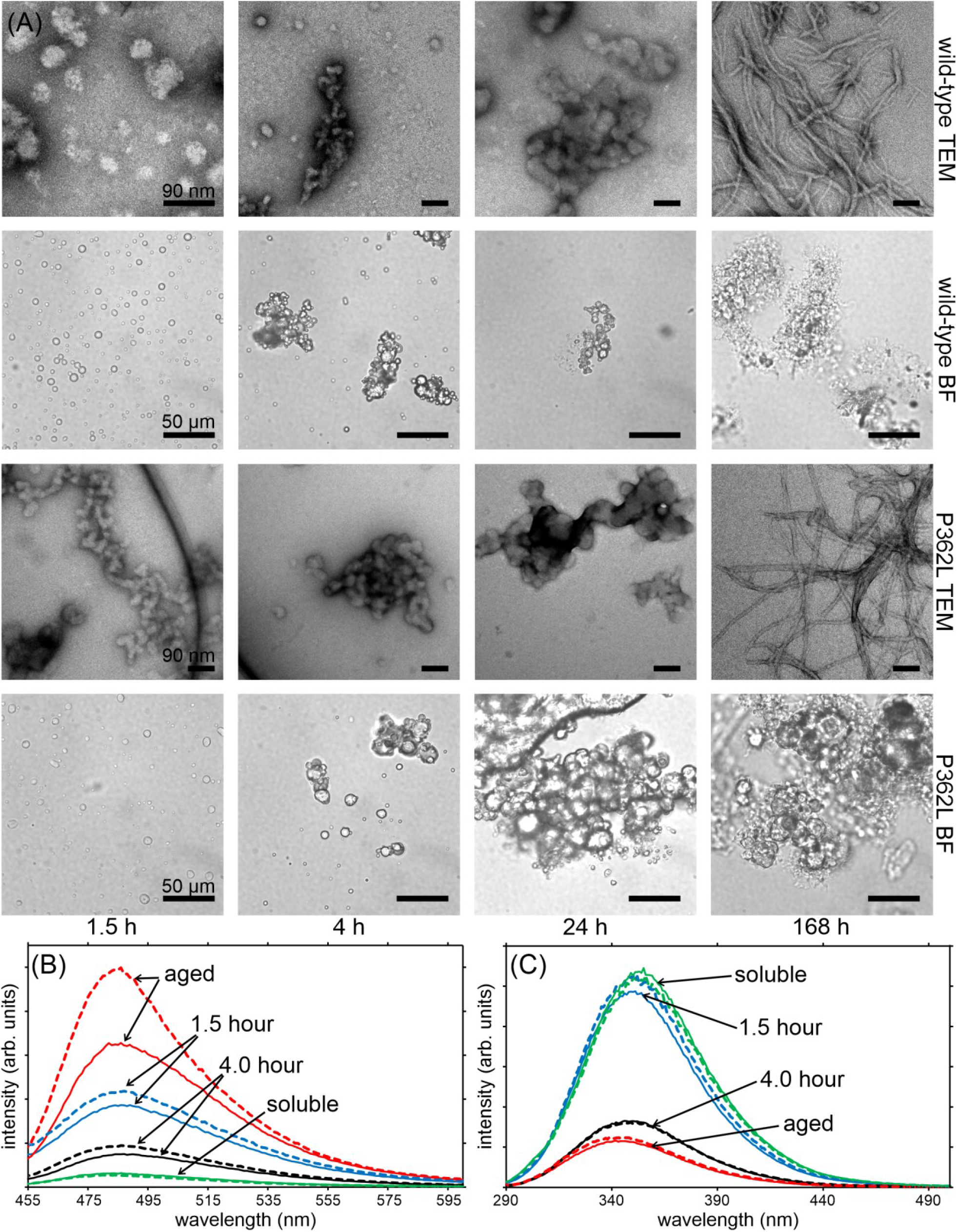
Droplet Aging of TIA1 LC Domain Wild-type and P362L Mutant Fibrils. (A) Bright field microscope images (BF) and negatively stained electron micrographs (TEM) of wild-type and P362L mutant TIA1 LC domain samples show the conversion of liquid droplets into amorphous aggregates that then convert into fibrils over the course of one week. (B) ThT fluorescence spectra recorded over the same time period show a slight increase in intensity after 1.5 h, which decreases at 4 h before increasing again over the course of a week. (C) Intrinsic Trp fluorescence spectra recorded over the same time period show a slight decrease in intensity and shift to lower wavelengths after 1.5 h, a trend that continues over the course of a week. In both (B) and (C), solid lines represent the data from wild type TIA1 LC domain sample and dashed lines represent data from P362L mutant TIA1 LC domain sample.

As the droplets are allowed to age over 24 h, they clump together. Figure 3A also shows TEM images recorded during the aging process, which reveal the formation of amorphous aggregates concomitantly with the disappearance of the liquid droplets in the bright field images. After one week without agitation, TEM images primarily show long, thin fibrils with variable degrees of bundling. The residual soluble protein concentrations for the wild-type and P362L liquid droplet preparations were 1.4 μM, indicating the aged liquid droplets have similar thermodynamic stabilities as the seeded fibrils.

The thioflavin T (ThT) fluorescence spectra in Figure 3B show an increase in fluorescence intensity for both wild-type and P362L mutant TIA1 LC domain in condensed phases when compared to denaturant solubilized protein. The intensity increases at 1.5 h, then decreases at 4 h before rising again at ∼1 week. An increase in ThT fluorescence suggests the presence of a cross-β conformation for TIA1 LC domain at 1.5 h and incubation times greater than 7 d, with reduced cross-β conformations at the intermediate 4 h time point. However, moderate increases in ThT fluorescence can arise from confinement of the fluorophore in a biomolecular condensate rather than the formation of rigid and extended cross-β structure.^32,34^ Both the wild-type and P362L mutant TIA1 LC domain follow a similar ThT time course, suggesting the liquid droplet to fibril transition is similar for both samples. The total increase in the ThT intensity is significantly larger for the P362L mutant sample than the wild-type sample and is consistent with previous ThT measurements on fibrils of full-length TIA1.^15^

The intrinsic tryptophan fluorescence spectra in Figure 3C show a steady decrease in intensity and a shift toward lower wavelengths over time for both wild-type and P362L mutant TIA1 LC domain. These spectra are consistent with one or more of the five Trp residues in the TIA1 LC domain losing accessibility to aqueous solvent.^35^ The large suppression of Trp fluorescence and distribution of the Trp residues in the TIA1 LC domain suggest that significant portions of the protein are protected from the solvent in the fibrils and to a lesser extent in the liquid droplets. The similar time-dependent Trp fluorescence signatures for both the wild-type and P362L mutant TIA1 LC domain indicate these processes are similar. The spectra at 4 h do not suggest an intermediate state with highly solvent exposed Trp residues as the liquid droplets age.

The ThT and Trp fluorescence measurements in Figures 3B and 3C are therefore consistent with both wild-type and P362L mutant TIA1 LC domains forming liquid droplets that transition into cross-β rich fibrils through an amorphous aggregate state on similar timescales.

### Wild-type and P362L Mutant TIA1 LC Domain Aged Droplets are Structurally Distinct from the Seeded Fibrils

Figure 4 shows cross-polarization based solid state NMR spectra of wild-type and P362L mutant TIA1 LC domain aged droplets. The carbon-carbon correlation spectra in Figures 4A and 4C show signal intensities that are consistent with ordered Thr, Ser, Pro, Ala, Asn or Asp, Gln or Glu, Phe, Tyr, Trp, Met, and Arg residues. A difference spectrum, where the spectrum of one sample is subtracted from that of another, highlights the most significant differences between the two spectra. Supplemental Figure S5 shows an overlay and difference spectrum for the wild-type and P362L mutant aged droplets, revealing these differences are minimal. Overall, these spectra appear highly similar to those of the seeded wild-type TIA1 LC domain fibrils in Figure 2A. However, the difference spectrum comparing the seeded fibril and aged droplet spectra for wild-type TIA1 LC in Figure 4E show that the sharp signals for residues W339–Q342, F344–N345, T347–S350, P352, M354, and P356–N357 are not present in the aged liquid droplet spectra. These spectra therefore indicate that the well-ordered conformation obtained from fibril seeding is largely not present in the aged liquid droplet samples. Overlays of the spectra used to calculate the difference spectrum in Figure 4E are shown in Supplemental Figure S5.

**Figure 4:**
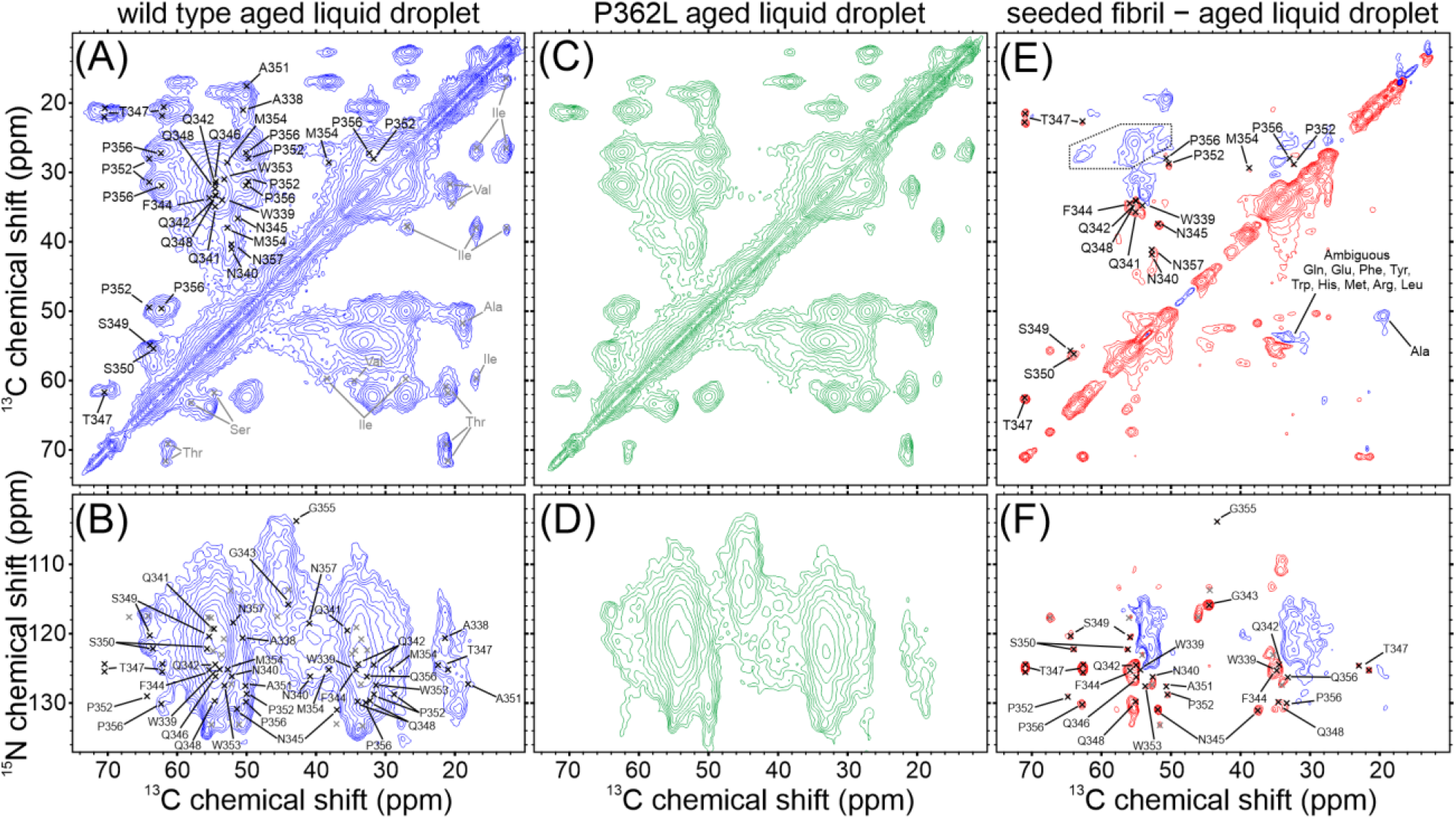
Solid State NMR Characterization of Wild-type and P362L Mutant TIA1 LC Domain Aged Liquid Droplets. (A) and (C) Carbon-carbon cross polarization-based correlation spectra of wild-type and P362L mutant TIA1 LC domain aged liquid droplets. (B) and (D) Nitrogen-carbon cross polarization-based correlation spectra of wild-type and P362L mutant TIA1 LC domain aged liquid droplets. (E) and (F) Carbon-carbon and nitrogen-carbon cross polarization based difference spectra obtained by subtracting a spectrum of the wild-type aged liquid droplets from a spectrum of the wild-type seeded fibrils. In the difference spectra, blue contours represent signals that are stronger in the wild-type aged liquid droplet sample and red contours represent signals that are stronger in the wild-type seeded fibril sample. The dashed outline in (E) indicates an experimental artifact in the spectrum. For all spectra, the black labels and marks are the unambiguously assigned signals from the seeded TIA1 LC domain fibrils, and the gray marks are the unassigned signals from the seeded TIA1 LC domain fibrils.

The nitrogen-carbon cross polarization-based spectra in Figures 4B and 4D for the wild-type and P362L mutant aged TIA1 LC domain droplets are also highly similar to one another. An overlay of these spectra, the difference spectra, and TEDOR-based spectra are provided in Supplemental Figure S5. Similar to the carbon-carbon spectra, the nitrogen-carbon difference spectrum for the wild-type seeded fibril and aged droplet spectra in Figure 4F reveals that the strong, sharp signals for residues W339, Q342–W353, and G355–P356 are not present in the spectrum of the aged droplets, indicating that the seeded fibril core structure is largely not present in the aged liquid droplet samples. Overlays of the spectra used to construct Figure 4F are shown in Supplemental Figure S5.

The spectra in Supplemental Figure S6 show that lowering the sample temperature for the aged droplet samples does not significantly change the appearance of the cross polarization and TEDOR based spectra. Reducing the temperature probes the presence of loosely ordered but conformationally homogenous structure in these samples.^30^ As molecular motions decrease with temperature, cross polarization and TEDOR based magnetization transfers should be more efficient due to stronger dipolar coupling interactions. Notably, the signals reporting on the seeded wild-type fibril conformation do not appear in these spectra. These low temperature data are therefore not consistent with a loosely ordered structure similar to the seeded wild-type conformation and further support that there are significant structural differences in molecular conformations between the seeded fibril and aged droplet TIA1 LC domain samples.

Scalar-based experiments were performed on the wild-type and P362L mutant TIA1 LC domain aged liquid droplets. No signals were observed in the spectra recorded under the same conditions as the seeded TIA1 LC domain fibrils. Lack of signal intensity in these spectra indicate that there are no residues with significant molecular motion in either aged liquid droplet sample. ^15^N and ^13^C relaxation measurements for these samples probe the presence of structural heterogeneity and molecular motion in the samples. ^15^N *T*_*2*_ values for the wild-type and P362L mutant aged droplets are 9.80 +/− 0.02 ms and 9.91 +/− 0.02 ms, respectively. The ^13^C *T*_*2*_ values for the wild-type and P362L mutant droplets are 2.92 +/− 0.01 ms and 2.93 +/− 0.01 ms, respectively. These measurements indicate that the intrinsic ^15^N and ^13^C linewidths (full width at half maximum) are on the order of 0.4 and 0.5 ppm. In the context of the 2D solid state NMR spectra shown in Figure 4, these relaxation parameters are consistent with the TIA1 LC domain monomers in the aged liquid droplets occupying an array of heterogeneous rigid conformations rather than a single conformation undergoing molecular motion.

## DISCUSSION

Here we have shown, (i) a seeded preparation of the TIA1 LC domain yields uniform fibrils (Figure 1), (ii) the seeded fibril core is formed by residues 338–357 in β-strand conformations (Figure 2, Supplemental Figures S1–S4), (iii) aging of TIA1 LC domain liquid droplets results in β-strand rich fibrils that are conformationally heterogeneous and structurally distinct from the seeded fibrils (Figures 3 and 4, Supplemental Figure 5), and (iv) the P362L mutation lies outside of the core forming region for the seeded fibrils and does not alter the structural characteristics of aged droplets (Figures 2 and 4, Supplemental Figures 5 and 6).

### Structure in Aged Liquid Droplets

Solid state NMR studies of condensate gelation are currently limited in number but are capable of providing highly informative atomic-resolution or residue-specific characterizations of molecular conformation and motion. Heterochromatin protein 1α (HP1α) was shown to undergo a droplet to gel transition characterized by increased uniform and rigid structure coexisting with highly mobile and disordered structure.^36^ However, the detailed conformation of the HP1α protein in the aged droplet sample was not characterized in the study. Aging of the fused in sarcoma (FUS) LC domain showed a similar increase of rigid structure in the presence of highly mobile and disordered regions of the protein.^37^ In this case, the aged droplets were predominantly composed of a single rigid molecular conformation with NMR chemical shifts indicative of β-strand structures, results that were supported by TEM images and ThT positive fluorescence measurements.^37^ Although the molecular conformation of the FUS LC domain in the aged droplets was similar to seeded FUS LC domain fibrils,^29^ significant differences in the solid state NMR spectra suggest the structural conversion was incomplete. Our recent study of aged droplets of the TAR DNA-binding protein 43 (TDP43) LC domain revealed the protein converges on a singular β-strand rich molecular conformation without seeding.^32^ In these aged liquid droplets however, there was evidence for conformational heterogeneity and only a small fraction of the LC domain exhibited rapid molecular motion.^32^

Our ThT fluorescence measurements show the TIA1 LC domain aged liquid droplets are rich in β-strand structure. However, the individual molecules in the droplets are conformationally heterogeneous given the lack of strong and sharp peaks in the solid state NMR spectra, in contrast to the more conformationally homogenous FUS and TDP43 LC domain aged droplets. Furthermore, significant portions of the TIA1 LC domain in both the seeded fibrils and aged droplets do not participate in the rigid core of the structure but have limited molecular motion and are likely loosely ordered, characteristics that differ from the HP1α protein and FUS LC domain but are similar to the TDP43 LC domain. We also note the surprising result that the ThT and Trp fluorescence spectra and TEM images of the TIA1 LC domain aging process suggest a non-fibrillar aggregate intermediate between the liquid droplet and β-strand rich fibril states, which was not observed for the HP1α, FUS-LC domain, or TDP43-LC domain proteins.^32,36,37^

Fibril seeding propagates specific fibril conformations for amyloid proteins like β-amyloid (Aβ), Tau, and α-synuclein.^38–40^ *In vitro*, seeding selects for the fibril conformation that has the fastest propagation rate, which may also be the most thermodynamically stable molecular conformation.^21^ However, different Aβ fibril polymorphs exhibit similar thermodynamic stabilities.^41^ While the TEM images and fluorescence data shown here indicate that the TIA1 LC domain droplets age into fibril structures visually similar to the seeded TIA1 LC domain, our solid state NMR measurements show the aged droplets exhibit a significant degree of conformational heterogeneity, contain a limited amount of the molecular conformation selected through seeding, and are inconsistent with the presence of another dominant conformation. These results suggest that the TIA1 LC domain is polymorphic regarding the formation of fibril structures, and our residual concentration measurements indicate that these conformations all have similar thermodynamic stabilities. Our observations suggest the intriguing possibility that the TIA1 LC domain conformations in aged droplets might be structurally compatible and could coexist within the same fibril.

### Fibril Forming Cores of LC Protein Domains

Residues A338–N357 of the TIA1 LC domain, comprised of two Trp, three Asn, four Gln, one Phe, one Thr, two Ser, two Pro, one Met, two Ala, and two Gly residues, have both differences and similarities with other fibril forming LC domain proteins. Here we discuss LC domains based on their known structural polymorphisms. For the monomorphic LC domains, both the FUS^29^ and heterogeneous ribonucleoprotein A2 (hnRNPA2)^11^ LC domain fibrils are formed by longer 57-residue sequences. Despite composing 16% of the TIA1 LC domain, there are only two Gly residues in the fibril core. The FUS LC domain structure contains 12 ordered Gly residues and the hnRNPA2 LC domain structure contains 20 ordered Gly residues. Furthermore, these structures contain a significant number of aromatic Phe and Tyr residues, eight for FUS and 11 for hnRNPA2, that likely participate in pi-pi interactions^42^ between adjacent monomers that stabilize the fibril conformations. The sequence identified here for the TIA1 LC domain only contains a single Phe residue but does have two Trp residues that may facilitate additional pi-pi interactions between adjacent monomers. Although there are weak signals we were unable to unambiguously assign in addition to those arising from the 20 residues in the rigid core of the seeded TIA1 LC domain fibrils, the 33 total signals observed are significantly fewer than the number of structured residues in the FUS and hnRNPA2 LC domain fibrils. Relative to the FUS and hnRNPA2 LC domain fibrils, the seeded TIA1 LC domain fibrils have significantly fewer ordered Gly residues, limited aromatic pi-pi interactions, and a shorter overall length.

There are common themes with respect to the monomorphic LC domain fibrils. First, Asn and Gln residues that can interact through sidechain hydrogen bonds with neighboring molecules to form polar zippers are prevalent. The FUS LC domain fibrils contain eight of these residues and the hnRNPA2 LC domain fibrils contain 14 of these residues, while the TIA1 LC domain fibril core contains seven of these residues. Second, ordered Pro residues, which are normally thought to disrupt secondary structure,^43^ are accommodated in these structures. TIA1 and hnRNPA2 contain two Pro residues, and FUS contains one Pro residue. The close backbone packing facilitated by LARK motifs^8^ in FUS and hnRNPA2 LC domain fibrils may facilitate ordering of Pro residues in the rigid fibril cores, and there is at least one possible LARK motif in the TIA1 LC domain fibril core region. Third, Thr and Ser residues can stabilize fibril structure in LC domains through intramolecular hydrogen bonding in lieu of an extended hydrophobic core.^29,44^ The FUS LC domain fibrils contain ten Ser and Thr residues positioned to form such hydrogen bond interactions, but the hnRNPA2 fibrils contain only a single pair of Ser in a similar arrangement. Although the TIA1 LC domain fibril core determined here only contains three Ser and Thr residues, this content is similar to that of the FUS LC fibrils when corrected for the total length of the fibril forming sequence of amino acids (16% for TIA1 LC domain and 18% for FUS LC).

Fibril formation in the context of purely pathological processes is typically associated with polymorphism.^45^ Hydrophobic amino acid content facilitating the formation of steric zippers is a distinguishing feature that separates reversible LC domain fibrillation and irreversible pathological fibril formation.^46^ The monomorphic FUS and hnRNPA2 LC domain fibrils contain limited buried hydrophobic residues. The TIA1 LC domain fibril core contains one Phe, two Pro, one Met, and two Ala which represent a larger percentage of the fibril core forming segment of the LC domain (25% for TIA1; 2% for FUS; and 7% for hnRNPA2). Trp and Tyr residues are excluded from this analysis due to the partial polar nature of these sidechains endowing the ability to form hydrogen bond interactions. Regarding purely hydrophobic residues, the TIA1 LC domain fibrils are more similar to those formed by the polymorphic TDP43 LC domain. Multiple fibril cores have been identified in the polymorphic TDP43 LC domain, all of which have significant hydrophobic residue content (23–44%).^32,47–49^ The higher prevalence of hydrophobic residues in the TIA1 LC domain fibril core may suggest that the propensity for liquid droplets to mature into polymorphic fibrils is likely to be more pathogenic than functional. However, the lack of TIA1 positive inclusions in patient tissues^15^ points to a more complicated role for TIA1 than purely forming pathogenic fibrils.

### TIA1 LC Domain Disease Mutations, Liquid Droplets, and Fibrils

According to the results of our measurements, the P362L mutation in the TIA1 LC domain resides outside the seeded fibril core and does not appreciably alter the molecular conformations present in the aged liquid droplets. In the full-length TIA1 protein, the primary effect of the Pro-to-Leu mutations and other disease associated LC domain mutations are an increased persistence time of stress-induced granules and more rapid increases in ThT fluorescence.^15,17^ The overall persistence time of the wild-type and P362L mutant droplets is shorter in our study of the TIA1 LC domain alone, but the rate of droplet assembly is similar to the full-length TIA1 protein.^15^ The increased rate and magnitude of ThT fluorescence for the P362L mutant are also consistent with the full length TIA1 protein.^15^ Pro-to-hydrophobic mutations in the TIA1 LC domain have been proposed to promote extension of β-strand structures.^17^ While our study does not directly address the structure of seeded TIA1 LC domain P362L mutant fibrils, our results clearly show that the seeded wild-type TIA1 LC domain fibrils can accommodate uniform and rigid conformations for Pro residues (P352 and P356). Furthermore, the heterogeneous molecular conformations we observe for the aged wild-type and P362L mutant droplets are not appreciably different. Given the similarities in maturation kinetics for our TIA1 LC domain droplets with the full-length TIA1 protein, we propose our data support that the P362L mutation mainly affects protein-protein interactions in the dynamic droplet state rather thermodynamically favoring a specific rigid fibrillar conformation. The increased persistence time and altered molecular interactions inside the droplets caused by mutations may therefore be responsible for nucleating aggregation of other proteins like TDP43. We note that considerable discussion has been devoted to the functional role of seeded TIA1 fibril formation.^20^ Our detailed characterization of the seeded fibril core may therefore provide a starting point for a more precise dissection of functional TIA1 assembly. Figure 5 depicts the two separate pathways for TIA1 LC domain assembly that are suggested by our results.

**Figure 5:**
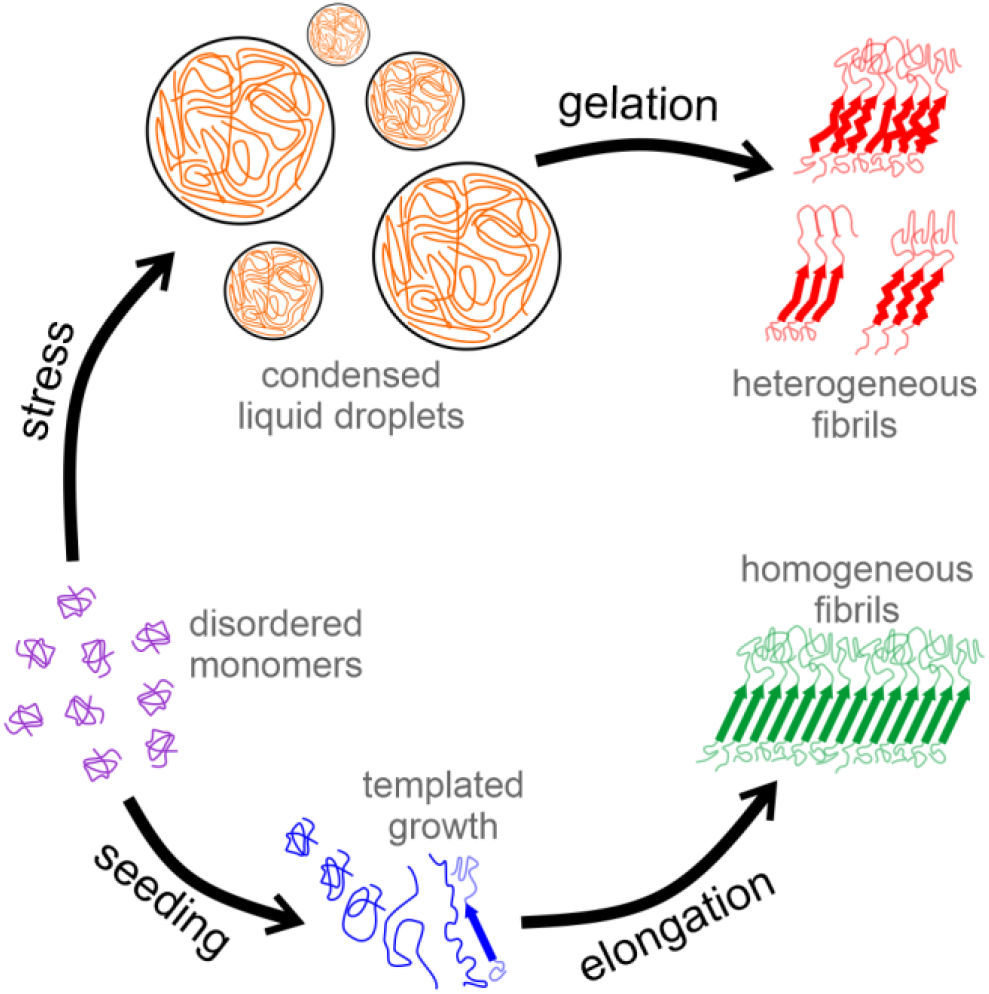
Possible Assembly Pathways for the TIA1 LC Domain. In one pathway, liquid droplets can form in response to stress stimuli. Disease mutations cause these structures to persist for abnormally long times and alter protein-protein interactions. The adoption of structurally heterogenous fibril conformations facilitates the pathological nucleation of other proteins. In a second pathway, seeds composed of molecules in well-defined conformations can template limited homogeneous fibril formation to bring proteins together functionally.

Characterizing TIA1 LC domain assembly is an important component of a broad search for understanding other functional and pathological activities governed by LC domain protein self-assembly. 30% of proteins in the human proteome have at least one LC domain.^8^ The amino acid biases of these proteins vary greatly. Guiding principles are emerging for the role of amino acid types and distributions for liquid droplet formation.^50,51^ However, functional amyloid-like fibril formation is also a key component of organizing cellular activity.^52^ The large number of missense mutations in LC domains linked to disease^14^ make it clear that the precise sequences of these proteins are important. Since functional LC domains differ in amino acid content from those of traditional and purely pathological amyloids, further characterizations of these fascinating and ubiquitous protein domains are needed.

## ACKNOWLEDGEMENTS

We wish to thank Dr. Steven L. McKnight for generously providing the TIA1 LC domain plasmid used in this work. Research reported in this publication was supported by the National Institute of General Medical Sciences of the National Institutes of Health under award number R35142892 to D.T.M. A portion of this work was performed at the National High Magnetic Field Laboratory, which is supported by the National Science Foundation Cooperative Agreement No. DMR-1644779 and the State of Florida. The instrumentation at the University of California, Davis, NMR facility is partly funded by the National Science Foundation under award number DBI-0722538. A portion of this work was supported through generous start up funding from the University of California, Davis to D.T.M. Dr. Jeff Walton provided support for the NMR instrumentation and Dr. Fei Guo provided support for the electron microscope at University of California, Davis Campus Core Facilities.

## AUTHOR CONTRIBUTIONS

Conceptualization, D.T.M.; Formal Analysis, Y.W. and D.T.M.; Investigation, Y.W., K.M.J., R.S., T.L., I.H., and D.T.M.; Writing — Original Draft, Y.W. and D.T.M.; Writing — Review & Editing, Y.W., I.H., and D.T.M.; Visualization, Y.W. and D.T.M.; Supervision, D.T.M.; Funding Acquisition, D.T.M.

## MATERIALS AND METHODS

### Protein Expression and Purification

His-tagged TIA1 LC domain (residues I291–Q386) wild type and P362L mutant proteins were recombinantly produced using a pHIS-parallel plasmid^53^ includes an N-terminal His-tag (MSYYHHHHHHDYDIPTTENLYFQGAMDPEF). The plasmid was transformed in BL21(DE3) *E. coli* cells. Cells were grown in a shaker incubator at 37 °C and 220 revolutions per minute (rpm). For unlabeled protein, bacterial cultures were grown in Luria broth media at 37 °C with shaking at 220 rpm in the presence of 100 mg/ml ampicillin to an optical density at 600 nm of 0.6–1.0 measured using a 1 cm pathlength cuvette before adding isopropyl β-D-thiogalactopyranoside (IPTG) to 0.5 mM to induce protein expression. The cultures were grown for another 3 h before harvesting the cells by centrifugation at 6,000 g for 15 min. The cell pellets were flash frozen in liquid nitrogen and stored at −80 °C until purification. For ^13^C and ^15^N labeled protein, bacterial cells were grown to an optical density at 600 nm of ∼2.0 in Dynamite Broth^54^. Cells from 2 L culture were harvested by centrifugation at 6,000 g for 10 min and transferred into 1 L of M9 minimal media using a serological pipette (45.8 mM Sodium phosphate dibasic heptahydrate, 22.0 mM Potassium phosphate monobasic, 8.6 mM sodium chloride, 2.0 mM magnesium chloride, 0.1 mM calcium chloride, and 100 μg/ml ampicillin 2.0 g of U-^13^C_6_ D-glucose, and 1.0 g of ^15^N ammonium chloride (Cambridge Isotope Labs)). Protein expression was induced after growing the cells for 0.5 h longer at 37 °C with 220 RPM shaking by adding IPTG to 0.5 mM. The cells were harvested 3 h later by centrifugation at 6,000 g for 15 min, flash frozen in liquid nitrogen, and stored at −80 °C until purification.

Purification procedures for the wild type and P362L mutant TIA1 LC domain were the same. Cell pellets were thawed on ice for ∼15 min and resuspended in 6 M guanidinium chloride, 50 mM tris(hydroxymethyl)aminomethane (Tris) pH 7.5, 500 mM sodium chloride, and 1% v/v Triton X-100 along with 3 pellets of EDTA-free Pierce Protease Inhibitor (ThermoFisher Scientific), and 0.25 mg/ml hen egg white lysozyme. The resuspended cells were sonicated using a Branson 250 Sonifier equipped with a 1/4 inch microtip in an ice-water bath for a total of 20 min at 30% output, in cycles of 0.3 s on and 3 s off, for a total of 1 min sonication. The lysed cells were centrifuged at 4 °C and 75,600 g for 30 min to remove insoluble material. TIA1 LC domain was isolated from the supernatant using a Bio-Rad NGC Discover 10 chromatography system and a 5 mL Ni-affinity column (Bio-Rad Bio-scale Mini Nuvia IMAC or GE Healthcare HisTrap FF crude). The column was equilibrated in 500 mM sodium chloride, 6 M urea, and 20 mM 4-(2-hydroxyethyl)-1-piperazineethanesulfonic acid (HEPES) pH 7.5; washed in 500 mM sodium chloride, 6 M urea, 20 mM imidazole, and 20 mM HEPES, pH 7.5; and eluted in a 500 mM sodium chloride, 6 M urea, and 20 mM HEPES, pH 7.5, using a 20–200 mM imidazole gradient. 1 mL aliquots of the elution peak containing TIA1 LC domain were prepared and stored at −80 °C. The presence and purity of TIA1 LC domain was confirmed by SDS-PAGE with Coomassie staining.

### TIA1 LC Domain P362L Site-directed Mutagenesis

The P362L mutant His-tagged TIA1 LC domain plasmid was prepared using forward and reverse primers containing the P362L mutation (5’- ccattttgcccttgaggcagttgcactccataatttg-3’ and 5’-caaattatggagtgcaactgcctcaagggcaaaatgg-3’, Integrated DNA Technologies). A polymerase chain reaction (PCR) was performed by mixing 0.5 μL of the pHis-parallel wild-type TIA1 LC domain plasmid (123.1 ng/μl), 2.5 μL of 10 μM forward and reverse primers, 5× Phusion GC buffer, 1 μL of 10 mM deoxynucleotides, 1.5 μL of dimethyl sulfoxide, and 0.5 μL of a Phusion DNA polymerase was mixed gently in a 50 μl volume. Polymerase, buffer and dNTPs were purchased from New England BioLabs. PCR was performed using a Bio-Rad T100 thermal cycler with an initial denaturation step at 98 °C for 30 s followed by 30 cycles of 7 s denaturation at 98 °C, 20 s of annealing at 72 °C, and extension for 2 min 42 s, and finally a final extension for 8 min at 72 °C. DpnI (New England Biolabs) was added to the PCR reaction vial and incubated at 37 °C for 1 h before storage at 4 °C. The PCR product was transformed into DH5α *E. coli* cells and grown in Luria broth media for 17 h at 37 °C with 220 RPM shaking. The amplified plasmid was purified using a Qiagen QIAprep Spin Miniprep kit. The sequence of the plasmid was confirmed by Sanger sequencing (Genewiz).

### Seeded TIA1 LC Domain Fibril Preparation

TIA1 LC domain seeds were prepared by concentrating the protein to ∼2.5 mg/mL, ∼6 mL using Millipore Amicon Ultra-15 3 kDa MWCO centrifugal filter and dialyzed (Spectrum Laboratories 1.7 ml/cm standard SpectraPor 1 RC Tubing) against 1 L of 20 mM HEPES, pH 7.5, buffer for 24 h to remove urea. The protein was harvested in 150 μL aliquots and flash frozen in liquid nitrogen and stored at −80 °C.

TIA1 LC domain fibrils were prepared by concentrating the protein to ∼2.5 mg/mL in a ∼6 mL volume using Millipore Amicon Ultra-15 3 kDa MWCO centrifugal filter and dialyzed (Spectrum Laboratories 1.7 ml/cm standard SpectraPor 1 RC Tubing) against 1 L of 20 mM HEPES, pH 7.5, buffer for 24 h to remove urea. The dialysis was harvested after ∼24 h. One aliquot of seeds stored at −80 °C was thawed to room temperature and diluted with 150 μL of 20 mM HEPES, pH 7.5, buffer. This was sonicated using a Branson 250 Sonifier equipped with a 1/8” microtip at 10% power in cycles of 0.1 s on and 1 s off for a total on time of 1 min. These seeds were then added to the harvested protein and left on benchtop for a week to quiescently form fibrils.

### TIA1 LC Domain Liquid Droplet Formation

For both wild type and P362L mutant TIA1 LC domain, a total of 15 mg purified protein was diluted to 1.2 mg/mL using 500 mM sodium chloride, 6 M urea, 200 mM imidazole, and 20 mM HEPES, pH 7.5 buffer and dialyzed against 1 L of 150 mM sodium chloride and 20 mM HEPES, pH 7.5, buffer. 65 μL aliquots were taken out of the dialysis tubing after gently pipetting to homogenize and mix the sample, at 90 min, 3 h, and 4 h for fluorescence measurements and microscopy imaging. The protein was transferred from the dialysis bag after 4 to a 50 mL conical tube and incubated on the benchtop at room temperature to form fibrils quiescently. 65 μL aliquots were also made after 24 h, 48 h, and 1 week after gently pipetting to homogenize the sample.

### Brightfield Microscopy

Liquid droplets were imaged on Olympus BX51 light microscope with differential interference contrast (DIC) filters, a 40× objective lens, and a Diagnostics Instruments RT Slider camera with 6-megapixel sampling. 3 μL of sample was dispensed onto a glass microscope slide and imaged immediately.

### ThT and Intrinsic Fluorescence Assays

A saturated ThT stock solution (>200 μM) was prepared by dissolving solid ThT (Acros Organics) in ultrapure water. The stock solution was sterile filtered using a syringe with a 0.22 μM PES membrane (Millipore). The ThT working concentration was 40 μM with 150 mM sodium chloride and 20 mM HEPES, pH 7.5. The ThT concentration was measured using absorbance at 412 nm and an extinction coefficient (ε412) of 36,000 M^−1^cm^−1^. All ThT solutions were stored at 4 °C, wrapped in foil, and used within 7 d. For ThT fluorescence measurements, 50 μL of the 40 μM ThT working solution was added directly into 50 μL of the protein sample and loaded into a glass quartz cuvette. For intrinsic fluorescence measurements, 50 μL of the protein solution was loaded directly into a glass quartz cuvette.

Intrinsic fluorescence and ThT fluorescence measurements were recorded on an Agilent Cary Eclipse fluorescence spectrophotometer and a 45 μL 10 mm × 10 mm quartz cuvette (Starna). The excitation and emission slit widths were 5 nm with a scan rate of 600 nm/min. The averaging time was 0.1 s with an interval of 1.0 nm. Intrinsic fluorescence data were recorded from 250 and 520 nm with an excitation wavelength of 280 nm. ThT data were acquired from 450 to 600 nm with an excitation wavelength at 440 nm.

### Transmission Electron Microscopy

Ted Pella UC-A lacey 400 mesh Cu grids were used for seeded TIA1 LC domain fibrils and Ted Pella ultrathin C film lacey carbon 300 mesh Au grids were used for both WT and P362L TIA1 LC domain liquid droplets. These were glow discharged before applying 5 μl of the TIA1 LC domain solutions incubating for 2 min. Bulk solution was blotted using a laboratory tissue and the grid was then washed twice by applying 5 μL of ultrapure water for ∼10 s and blotting away the water between each application. The grid was stained using 5 μL of 3% (w/v) uranyl acetate, incubated for 10 s, blotted with a tissue, and air-dried. Images were obtained on a JEOL-1230 electron microscope equipped with a 2K × 2K Tietz CCD camera operating at 100 keV.

### Solid State NMR Data Collection

Seeded TIA1 LC domain fibrils were pelleted at 30,000 g at 25 °C for 30 min. 10–15 mg of seeded TIA1 LC domain hydrated pellets were transferred into 3.2 mm thin-walled pencil-style zirconia rotors (Revolution NMR) with a spatula and packed by centrifugation at 25,000 g for ∼30 h at 8 °C. TIA1 LC domain wild-type and mutant aged liquid droplet fibrils were pelleted at 233,000 g at 12 °C overnight (∼20 h). 10–15 mg of TIA1 LC domain hydrated pellets were transferred into 3.2 mm thin-walled pencil-style zirconia rotors (Revolution NMR) with a spatula and packed by centrifugation at 25,000 g for 20 min at 12 °C. The rotor drive tip and top cap were sealed with cyanoacrylate gel. Residual soluble protein concentration was determined immediately after centrifugation by absorbance measurements at 280 nm using an Eppendorf Biospectrometer Kinetic, a μCuvette with a 1 mm pathlength, and an extinction coefficient calculated from the protein primary sequence.

Experiments were performed on 18.8 T magnets at the UC Davis NMR Campus Core Facility (Davis, CA) and the National High Magnetic Field Laboratory (Tallahassee, FL). BlackFox NMR and Low-E triple resonance 3.2 mm MAS probes were used for all experiments. A table of data acquisition parameters is provided in Supplementary Table S1. Unless otherwise specified, the sample temperature was ∼10 °C. The observed chemical shifts were externally referenced to the DSS scale with the ^13^C downfield peak of Adamantane at 40.48 ppm using a dehydrated sample of 1-^13^C labeled Ala powder. ^13^C *T*_*2*_ measurements were made by inserting a variable length spin-echo period immediately after the cross polarization step in a ^1^H-^13^C 1D experiment. ^15^N *T*_*2*_ measurements were made by inserting a variable length spin-echo period between the ^1^H-^15^N and ^15^N-^13^CA cross polarization steps in a 1D NCA experiment. Echo periods were varied up to 10.2 ms for ^13^C and 15.2 ms for ^15^N and utilized high power ^1^H decoupling. The integrated non-glycyl signal intensity was fit to single exponentials in TopSpin 3.6 software. For all spectra, carbon-carbon correlation spectra are plotted with contours increasing by a factor of 1.4. Nitrogen-carbon correlation spectra are plotted with contours increasing by a factor of 1.25.

### Residue-Specific Assignments

Chemical shift peak tables representing NCACX, NCOCX, and CANCO chemical shift correlations were constructed from the 3D CP-based datasets^25^ recorded on the seeded TIA1 LC domain fibril sample. A 2D TEDOR-NCACX experiment^26^ was used to identify the NCACX signals for proline residues. Signal assignment was achieved using the MCASSSIGN algorithm^28^ with a procedure that generally followed our previous assignments for the LC domains of Tm1^30^, hnRNPA2^31^, FUS^29^, and TDP43^32^. The input signal tables are presented in Supplemental Tables S2–S4. Due to significant overlap in the CB/CG region corresponding to residues such as Arg, His, Ile, Gln, Glu, Met, Trp, Tyr, Phe, and Leu, signals in this region were given generous uncertainties and allowed to be assigned to multiple residues in initial calculations. The MCASSIGN algorithm was run twice, each round consisting of 50 independent calculations with 20 steps and 10^7^ iterations per step. The GOOD, BAD, EDGE, and USED weights were ramped from (0–10), (10–60), (0–8), (0–6) during the annealing calculation. After the first round of calculations, signals were assigned for residues T347–N357 in 100% of the calculations. These assignments were fixed for a second round of calculations, which resulted in the same signals being assigned to residues A338–Q346 in 100% of the calculations. Subsequent calculations failed to converge on a unique result due to the large CB/CG uncertainties assigned for some signals and the repetitive nature of the TIA1 LC domain sequence. The calculation was run with the only His-tag as the input sequence resulting in no signals being assigned consistently to any stretch of residues greater than three, ruling out the possibility of the His-tag being significantly ordered in the fibril structure. A calculation run such that signals could not be assigned to residues A338–N357 resulted in no significant assignment of signals outside this region, ruling out the possibility that some of the chemical shifts derived from multiple residues in similar chemical environments and the possibility of a longer stretch of the TIA1 LC domain immobilized in the fibril core. Finally, a calculation run on the complete TIA1 LC domain with the NCOCX signals assigned to A338–N357 omitted but complete signal tables for the NCACX and CANCO data did not result in any significant assignments, ruling out the possibility of two different strongly ordered conformations for residues A338–N357. There are two sets of observed Thr signals that only differ significantly in their CG2 chemical shifts. The assignment process was unable to distinguish these sites due to their identical CA and CB chemical shifts and nearly identical amide N chemical shifts. Importantly, neither of these signals were ever consistently assigned to the very C-terminal Thr site or the two Thr sites in the His-tag.

## REFERENCES

(1) Standart, N.; Weil, D. P-Bodies: Cytosolic Droplets for Coordinated MRNA Storage. Trends in Genetics 2018, 34 (8), 612–626. https://doi.org/10.1016/j.tig.2018.05.005.

(2) Knowles, R. B.; Sabry, J. H.; Martone, M. E.; Deerinck, T. J.; Ellisman, M. H.; Bassell, G. J.; Kosik, K. S. Translocation of RNA Granules in Living Neurons. J. Neurosci. 1996, 16 (24), 7812–7820. https://doi.org/10.1523/JNEUROSCI.16-24-07812.1996.

(3) Mittag, T.; Parker, R. Multiple Modes of Protein–Protein Interactions Promote RNP Granule Assembly. Journal of Molecular Biology 2018, 430 (23), 4636–4649. https://doi.org/10.1016/j.jmb.2018.08.005.

(4) Molliex, A.; Temirov, J.; Lee, J.; Coughlin, M.; Kanagaraj, A. P.; Kim, H. J.; Mittag, T.; Taylor, J. P. Phase Separation by Low Complexity Domains Promotes Stress Granule Assembly and Drives Pathological Fibrillization. Cell 2015, 163 (1), 123–133. https://doi.org/10.1016/j.cell.2015.09.015.

(5) Corbet, G. A.; Parker, R. RNP Granule Formation: Lessons from P-Bodies and Stress Granules. Cold Spring Harb Symp Quant Biol 2019, 84, 203–215. https://doi.org/10.1101/sqb.2019.84.040329.

(6) Banani, S. F.; Lee, H. O.; Hyman, A. A.; Rosen, M. K. Biomolecular Condensates: Organizers of Cellular Biochemistry. Nature Reviews Molecular Cell Biology 2017, 18 (5), 285–298. https://doi.org/10.1038/nrm.2017.7.

(7) Kumari, B.; Kumar, R.; Kumar, M. Low Complexity and Disordered Regions of Proteins Have Different Structural and Amino Acid Preferences. Mol. BioSyst. 2015, 11 (2), 585–594. https://doi.org/10.1039/C4MB00425F.

(8) Hughes, M. P.; Sawaya, M. R.; Boyer, D. R.; Goldschmidt, L.; Rodriguez, J. A.; Cascio, D.; Chong, L.; Gonen, T.; Eisenberg, D. S. Atomic Structures of Low-Complexity Protein Segments Reveal Kinked β Sheets That Assemble Networks. Science 2018, 359 (6376), 698–701. https://doi.org/10.1126/science.aan6398.

(9) Fomicheva, A.; Ross, E. D. From Prions to Stress Granules: Defining the Compositional Features of Prion-Like Domains That Promote Different Types of Assemblies. International Journal of Molecular Sciences 2021, 22 (3), 1251. https://doi.org/10.3390/ijms22031251.

(10) Kato, M.; McKnight, S. L. Cross-β Polymerization of Low Complexity Sequence Domains. Cold Spring Harb Perspect Biol 2017, 9 (3), a023598. https://doi.org/10.1101/cshperspect.a023598.

(11) Lu, J.; Cao, Q.; Hughes, M. P.; Sawaya, M. R.; Boyer, D. R.; Cascio, D.; Eisenberg, D. S. CryoEM Structure of the Low-Complexity Domain of HnRNPA2 and Its Conversion to Pathogenic Amyloid. Nature Communications 2020, 11 (1), 4090. https://doi.org/10.1038/s41467-020-17905-y.

(12) Taylor, J. P.; Brown, R. H.; Cleveland, D. W. Decoding ALS: From Genes to Mechanism. Nature 2016, 539 (7628), 197–206. https://doi.org/10.1038/nature20413.

(13) Finger, E. C. Frontotemporal Dementias. Continuum (Minneap Minn) 2016, 22 (2 Dementia), 464–489. https://doi.org/10.1212/CON.0000000000000300.

(14) Harrison, A. F.; Shorter, J. RNA-Binding Proteins with Prion-like Domains in Health and Disease. Biochemical Journal 2017, 474 (8), 1417–1438. https://doi.org/10.1042/BCJ20160499.

(15) Mackenzie, I. R.; Nicholson, A. M.; Sarkar, M.; Messing, J.; Purice, M. D.; Pottier, C.; Annu, K.; Baker, M.; Perkerson, R. B.; Kurti, A.; Matchett, B. J.; Mittag, T.; Temirov, J.; Hsiung, G.-Y. R.; Krieger, C.; Murray, M. E.; Kato, M.; Fryer, J. D.; Petrucelli, L.; Zinman, L.; Weintraub, S.; Mesulam, M.; Keith, J.; Zivkovic, S. A.; Hirsch-Reinshagen, V.; Roos, R. P.; Zuchner, S.; Graff-Radford, N. R.; Petersen, R. C.; Caselli, R. J.; Wszolek, Z. K.; Finger, E.; Lippa, C.; Lacomis, D.; Stewart, H.; Dickson, D. W.; Kim, H. J.; Rogaeva, E.; Bigio, E.; Boylan, K. B.; Taylor, J. P.; Rademakers, R. TIA1 Mutations in Amyotrophic Lateral Sclerosis and Frontotemporal Dementia Promote Phase Separation and Alter Stress Granule Dynamics. Neuron 2017, 95 (4), 808–816.e9. https://doi.org/10.1016/j.neuron.2017.07.025.

(16) Wang, I.; Hennig, J.; Jagtap, P. K. A.; Sonntag, M.; Valcárcel, J.; Sattler, M. Structure, Dynamics and RNA Binding of the Multi-Domain Splicing Factor TIA-1. Nucleic Acids Research 2014, 42 (9), 5949–5966. https://doi.org/10.1093/nar/gku193.

(17) Ding, X.; Gu, S.; Xue, S.; Luo, S.-Z. Disease-Associated Mutations Affect TIA1 Phase Separation and Aggregation in a Proline-Dependent Manner. Brain Research 2021, 1768, 147589. https://doi.org/10.1016/j.brainres.2021.147589.

(18) Fritzsching, K. J.; Yang, Y.; Pogue, E. M.; Rayman, J. B.; Kandel, E. R.; McDermott, A. E. Micellar TIA1 with Folded RNA Binding Domains as a Model for Reversible Stress Granule Formation. PNAS 2020, 117 (50), 31832–31837. https://doi.org/10.1073/pnas.2007423117.

(19) Kato, M.; Han, T. W.; Xie, S.; Shi, K.; Du, X.; Wu, L. C.; Mirzaei, H.; Goldsmith, E. J.; Longgood, J.; Pei, J.; Grishin, N. V.; Frantz, D. E.; Schneider, J. W.; Chen, S.; Li, L.; Sawaya, M. R.; Eisenberg, D.; Tycko, R.; McKnight, S. L. Cell-Free Formation of RNA Granules: Low Complexity Sequence Domains Form Dynamic Fibers within Hydrogels. Cell 2012, 149 (4), 753–767. https://doi.org/10.1016/j.cell.2012.04.017.

(20) Rayman, J. B.; Kandel, E. R. TIA-1 Is a Functional Prion-Like Protein. Cold Spring Harb Perspect Biol 2017, 9 (5), a030718. https://doi.org/10.1101/cshperspect.a030718.

(21) Qiang, W.; Yau, W.-M.; Tycko, R. Structural Evolution of Iowa Mutant β-Amyloid Fibrils from Polymorphic to Homogeneous States under Repeated Seeded Growth. J. Am. Chem. Soc. 2011, 133 (11), 4018–4029. https://doi.org/10.1021/ja109679q.

(22) O’Nuallain, B.; Shivaprasad, S.; Kheterpal, I.; Wetzel, R. Thermodynamics of Aβ(1-40) Amyloid Fibril Elongation. Biochemistry 2005, 44 (38), 12709–12718. https://doi.org/10.1021/bi050927h.

(23) Takegoshi, K.; Nakamura, S.; Terao, T. 13C–1H Dipolar-Driven 13C–13C Recoupling without 13C Rf Irradiation in Nuclear Magnetic Resonance of Rotating Solids. J. Chem. Phys. 2003, 118 (5), 2325–2341. https://doi.org/10.1063/1.1534105.

(24) Takegoshi, K.; Nakamura, S.; Terao, T. 13C–1H Dipolar-Assisted Rotational Resonance in Magic-Angle Spinning NMR. Chemical Physics Letters 2001, 344 (5), 631–637. https://doi.org/10.1016/S0009-2614(01)00791-6.

(25) Franks, W. T.; Zhou, D. H.; Wylie, B. J.; Money, B. G.; Graesser, D. T.; Frericks, H. L.; Sahota, G.; Rienstra, C. M. Magic-Angle Spinning Solid-State NMR Spectroscopy of the ?1 Immunoglobulin Binding Domain of Protein G (GB1): 15N and 13C Chemical Shift Assignments and Conformational Analysis. J. Am. Chem. Soc. 2005, 127 (35), 12291–12305. https://doi.org/10.1021/ja044497e.

(26) Jaroniec, C. P.; Filip, C.; Griffin, R. G. 3D TEDOR NMR Experiments for the Simultaneous Measurement of Multiple Carbon-Nitrogen Distances in Uniformly 13C,15N-Labeled Solids. J. Am. Chem. Soc. 2002, 124 (36), 10728–10742. https://doi.org/10.1021/ja026385y.

(27) Morris, G. A.; Freeman, R. Enhancement of Nuclear Magnetic Resonance Signals by Polarization Transfer. J. Am. Chem. Soc. 1979, 101 (3), 760–762. https://doi.org/10.1021/ja00497a058.

(28) Tycko, R.; Hu, K.-N. A Monte Carlo/Simulated Annealing Algorithm for Sequential Resonance Assignment in Solid State NMR of Uniformly Labeled Proteins with Magic-Angle Spinning. Journal of Magnetic Resonance 2010, 205 (2), 304–314. https://doi.org/10.1016/j.jmr.2010.05.013.

(29) Murray, D. T.; Kato, M.; Lin, Y.; Thurber, K. R.; Hung, I.; McKnight, S. L.; Tycko, R. Structure of FUS Protein Fibrils and Its Relevance to Self-Assembly and Phase Separation of Low-Complexity Domains. Cell 2017, 171 (3), 615–627.e16. https://doi.org/10.1016/j.cell.2017.08.048.

(30) Sysoev, V. O.; Kato, M.; Sutherland, L.; Hu, R.; McKnight, S. L.; Murray, D. T. Dynamic Structural Order of a Low-Complexity Domain Facilitates Assembly of Intermediate Filaments. PNAS 2020, 117 (38), 23510–23518. https://doi.org/10.1073/pnas.2010000117.

(31) Murray, D. T.; Zhou, X.; Kato, M.; Xiang, S.; Tycko, R.; McKnight, S. L. Structural Characterization of the D290V Mutation Site in HnRNPA2 Low-Complexity–Domain Polymers. PNAS 2018, 115 (42), E9782–E9791. https://doi.org/10.1073/pnas.1806174115.

(32) Fonda, B. D.; Jami, K. M.; Boulos, N. R.; Murray, D. T. Identification of the Rigid Core for Aged Liquid Droplets of an RNA-Binding Protein Low Complexity Domain. J. Am. Chem. Soc. 2021, 143 (17), 6657–6668. https://doi.org/10.1021/jacs.1c02424.

(33) Shen, Y.; Bax, A. Protein Structural Information Derived from NMR Chemical Shift with the Neural Network Program TALOS-N. In Artificial Neural Networks; Cartwright, H., Ed.; Methods in Molecular Biology; Springer: New York, NY, 2015; pp 17–32. https://doi.org/10.1007/978-1-4939-2239-0_2.

(34) Friedhoff, P.; Schneider, A.; Mandelkow, E.-M.; Mandelkow, E. Rapid Assembly of Alzheimer-like Paired Helical Filaments from Microtubule-Associated Protein Tau Monitored by Fluorescence in Solution. Biochemistry 1998, 37 (28), 10223–10230. https://doi.org/10.1021/bi980537d.

(35) Royer, C. A. Probing Protein Folding and Conformational Transitions with Fluorescence. Chem. Rev. 2006, 106 (5), 1769–1784. https://doi.org/10.1021/cr0404390.

(36) Ackermann, B. E.; Debelouchina, G. T. Heterochromatin Protein HP1α Gelation Dynamics Revealed by Solid-State NMR Spectroscopy. Angewandte Chemie International Edition 2019, 58 (19), 6300–6305. https://doi.org/10.1002/anie.201901141.

(37) Berkeley, R. F.; Kashefi, M.; Debelouchina, G. T. Real-Time Observation of Structure and Dynamics during the Liquid-to-Solid Transition of FUS LC. Biophysical Journal 2021, 120 (7), 1276–1287. https://doi.org/10.1016/j.bpj.2021.02.008.

(38) Frost, B.; Ollesch, J.; Wille, H.; Diamond, M. I. Conformational Diversity of Wild-Type Tau Fibrils Specified by Templated Conformation Change. J Biol Chem 2009, 284 (6), 3546–3551. https://doi.org/10.1074/jbc.M805627200.

(39) Luk, K. C.; Song, C.; O’Brien, P.; Stieber, A.; Branch, J. R.; Brunden, K. R.; Trojanowski, J. Q.; Lee, V. M.-Y. Exogenous α-Synuclein Fibrils Seed the Formation of Lewy Body-like Intracellular Inclusions in Cultured Cells. Proceedings of the National Academy of Sciences 2009, 106 (47), 20051–20056. https://doi.org/10.1073/pnas.0908005106.

(40) Petkova, A. T.; Leapman, R. D.; Guo, Z.; Yau, W.-M.; Mattson, M. P.; Tycko, R. Self-Propagating, Molecular-Level Polymorphism in Alzheimer’s ß-Amyloid Fibrils. Science 2005, 307 (5707), 262–265. https://doi.org/10.1126/science.1105850.

(41) Qiang, W.; Kelley, K.; Tycko, R. Polymorph-Specific Kinetics and Thermodynamics of β-Amyloid Fibril Growth. J. Am. Chem. Soc. 2013, 135 (18), 6860–6871. https://doi.org/10.1021/ja311963f.

(42) McGaughey, G. B.; Gagné, M.; Rappé, A. K. π-Stacking Interactions: ALIVE AND WELL IN PROTEINS *. Journal of Biological Chemistry 1998, 273 (25), 15458–15463. https://doi.org/10.1074/jbc.273.25.15458.

(43) Morgan, A. A.; Rubenstein, E. Proline: The Distribution, Frequency, Positioning, and Common Functional Roles of Proline and Polyproline Sequences in the Human Proteome. PLOS ONE 2013, 8 (1), e53785. https://doi.org/10.1371/journal.pone.0053785.

(44) Murray, D. T.; Tycko, R. Side Chain Hydrogen-Bonding Interactions within Amyloid-like Fibrils Formed by the Low-Complexity Domain of FUS: Evidence from Solid State Nuclear Magnetic Resonance Spectroscopy. Biochemistry 2020, 59 (4), 364–378. https://doi.org/10.1021/acs.biochem.9b00892.

(45) Tycko, R. Amyloid Polymorphism: Structural Basis and Neurobiological Relevance. Neuron 2015, 86 (3), 632–645. https://doi.org/10.1016/j.neuron.2015.03.017.

(46) Murray, K. A.; Evans, D.; Hughes, M. P.; Sawaya, M. R.; Hu, C. J.; Houk, K. N.; Eisenberg, D. Extended β-Strands Contribute to Reversible Amyloid Formation. ACS Nano 2022, 16 (2), 2154–2163. https://doi.org/10.1021/acsnano.1c08043.

(47) Zhuo, X.-F.; Wang, J.; Zhang, J.; Jiang, L.-L.; Hu, H.-Y.; Lu, J.-X. Solid-State NMR Reveals the Structural Transformation of the TDP-43 Amyloidogenic Region upon Fibrillation. J. Am. Chem. Soc. 2020, 142 (7), 3412–3421. https://doi.org/10.1021/jacs.9b10736.

(48) Li, Q.; Babinchak, W. M.; Surewicz, W. K. Cryo-EM Structure of Amyloid Fibrils Formed by the Entire Low Complexity Domain of TDP-43. Nature Communications 2021, 12 (1), 1620. https://doi.org/10.1038/s41467-021-21912-y.

(49) Arseni, D.; Hasegawa, M.; Murzin, A. G.; Kametani, F.; Arai, M.; Yoshida, M.; Ryskeldi-Falcon, B. Structure of Pathological TDP-43 Filaments from ALS with FTLD. Nature 2022, 601 (7891), 139–143. https://doi.org/10.1038/s41586-021-04199-3.

(50) Bremer, A.; Farag, M.; Borcherds, W. M.; Peran, I.; Martin, E. W.; Pappu, R. V.; Mittag, T. Deciphering How Naturally Occurring Sequence Features Impact the Phase Behaviours of Disordered Prion-like Domains. Nat. Chem. 2022, 14 (2), 196–207. https://doi.org/10.1038/s41557-021-00840-w.

(51) Martin, E. W.; Holehouse, A. S.; Peran, I.; Farag, M.; Incicco, J. J.; Bremer, A.; Grace, C. R.; Soranno, A.; Pappu, R. V.; Mittag, T. Valence and Patterning of Aromatic Residues Determine the Phase Behavior of Prion-like Domains. Science 2020, 367 (6478), 694–699. https://doi.org/10.1126/science.aaw8653.

(52) Sanders, D. W.; Kaufman, S. K.; Holmes, B. B.; Diamond, M. I. Prions and Protein Assemblies That Convey Biological Information in Health and Disease. Neuron 2016, 89 (3), 433–448. https://doi.org/10.1016/j.neuron.2016.01.026.

(53) Sheffield, P.; Garrard, S.; Derewenda, Z. Overcoming Expression and Purification Problems of RhoGDI Using a Family of “Parallel” Expression Vectors. Protein Expression and Purification 1999, 15 (1), 34–39. https://doi.org/10.1006/prep.1998.1003.

(54) Taylor, T.; Denson, J.-P.; Esposito, D. Optimizing Expression and Solubility of Proteins in E. Coli Using Modified Media and Induction Parameters. In Heterologous Gene Expression in E.coli: Methods and Protocols; Burgess-Brown, N. A., Ed.; Methods in Molecular Biology; Springer: New York, NY, 2017; pp 65–82. https://doi.org/10.1007/978-1-4939-6887-9_5.

