## Supplemental Figures and Tables for "Liquid droplet aging and fibril formation of the stress granule protein TIA1 low complexity domain"

**Supplemental Video 1: TIA1 LC Domain Wild-type Droplet Dynamics Related to Figure 3.** Wild-type TIA1-LC liquid droplets after 1.5 h of dialysis. Each frame is taken 1 s apart and the video covers 50 s of real time.

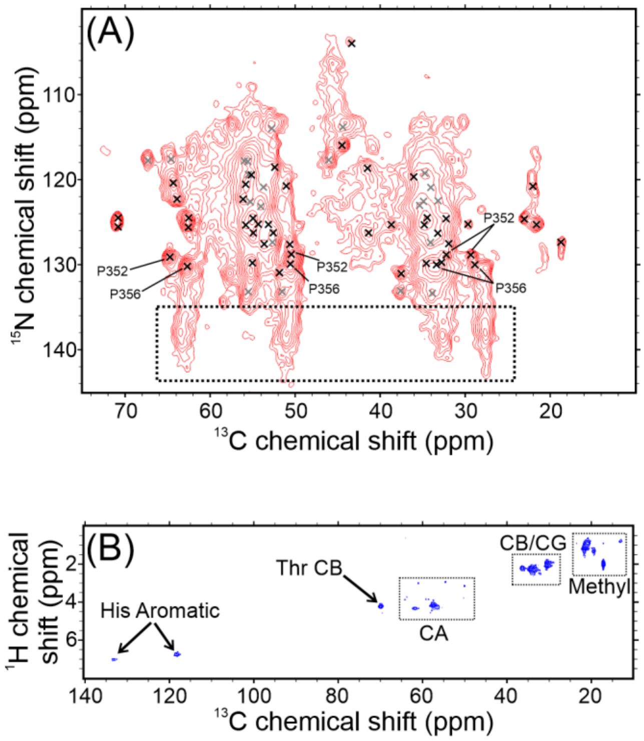


**Supplemental Figure 1: Additional 2D Solid State NMR Spectra of the Seeded TIA1 LC Domain Fibrils Related to Figure 2**. (A) cross polarization-based 2D NCACX-TEDOR spectrum of the wild-type seeded TIA1-LC fibrils highlights ordered Pro residues. Pro residues with rigid homogeneous conformations are labeled and Pro residues with rigid but heterogeneous conformations are indicated with a dashed box. Black marks are the unambiguously assigned signals for the wild type seeded TIA1 LC domain fibrils. Gray marks represent signals from the wild type seeded TIA1 LC domain fibrils that were not unambiguously assigned in this work. (B) Scalar-based 2D $^{1}$H-$^{13}$C INEPT spectrum of the wild-type seeded TIA1 LC fibrils highlights highly mobile sites. Signals that can be unambiguously assigned to an amino acid type are two aromatic His sidechain sites and a Thr CB site. Other signals are observed from CA, CB or CG, and methyl groups in amino acid sidechains. The contours are drawn at intensity values increasing by a factor of 1.25 in (A) and 1.5 in (B).

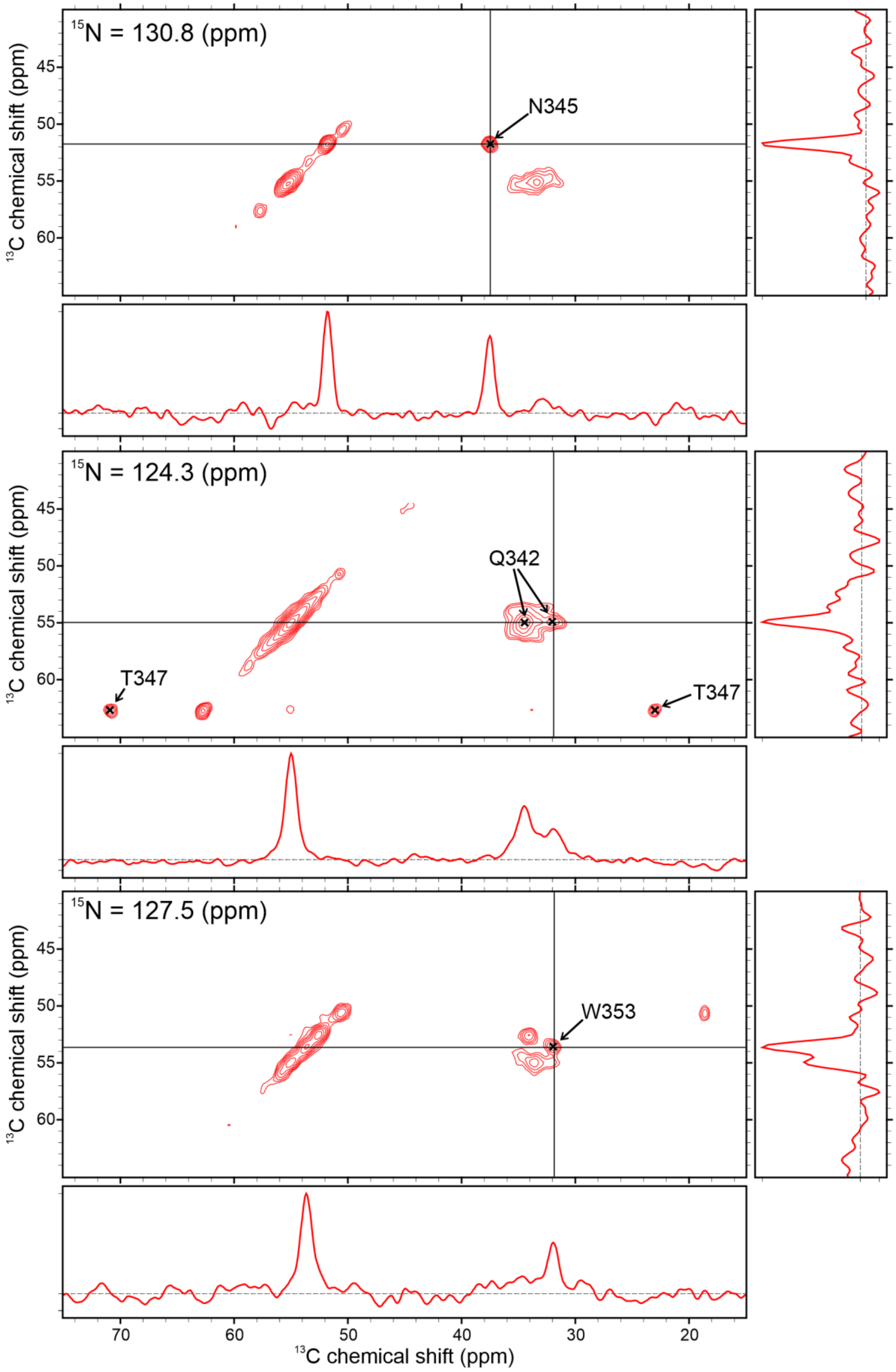

15N = 130.8 (ppm)
N345
15N = 124.3 (ppm)
Q342
T347
T347
15N = 127.5 (ppm)
W353
13C chemical shift (ppm)
13C chemical shift (ppm)
13C chemical shift (ppm)
13C chemical shift (ppm)

**Supplemental Figure 2: 2D Planes From the 3D NCACX Spectrum of the Wild-type Seeded TIA1 LC Domain Fibrils Related to Figure 2.** These 2D planes demonstrate the signal to noise and increased resolution in the 3D cross polarization-based NCACX experiment. The side and bottom panels next to each plane contain slices from the data at the location of the horizontal and vertical lines. The contours are drawn at intensity values increasing by a factor of 1.3.

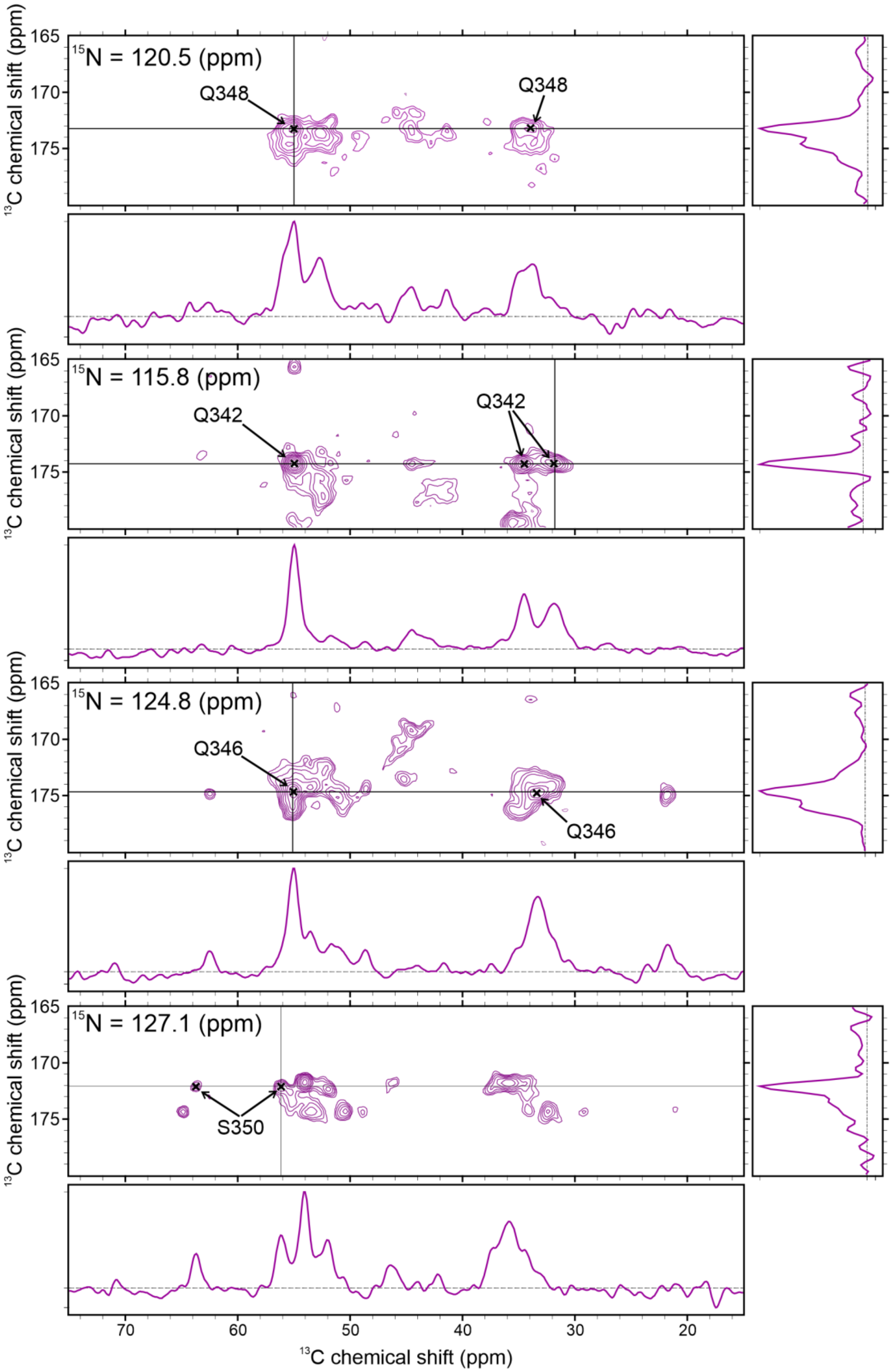

$^{15}$N = 120.5 (ppm)
Q348
Q348
$^{15}$N = 115.8 (ppm)
Q342
Q342
$^{15}$N = 124.8 (ppm)
Q346
Q346
$^{15}$N = 127.1 (ppm)
S350
$^{13}$C chemical shift (ppm)
$^{13}$C chemical shift (ppm)
165
170
175
70
60
50
40
30
20

**Supplemental Figure 3: 2D Planes From the 3D NCOCX Spectrum of the Wild-type Seeded TIA1 LC Domain Fibrils Related to Figure 2.** These 2D planes demonstrate the signal to noise and increased resolution in the 3D cross polarization-based NCOCX experiment. The side and bottom panels next to each plane contain slices from the data at the location of the horizontal and vertical lines. The contours are drawn at intensity values increasing by a factor of 1.3.

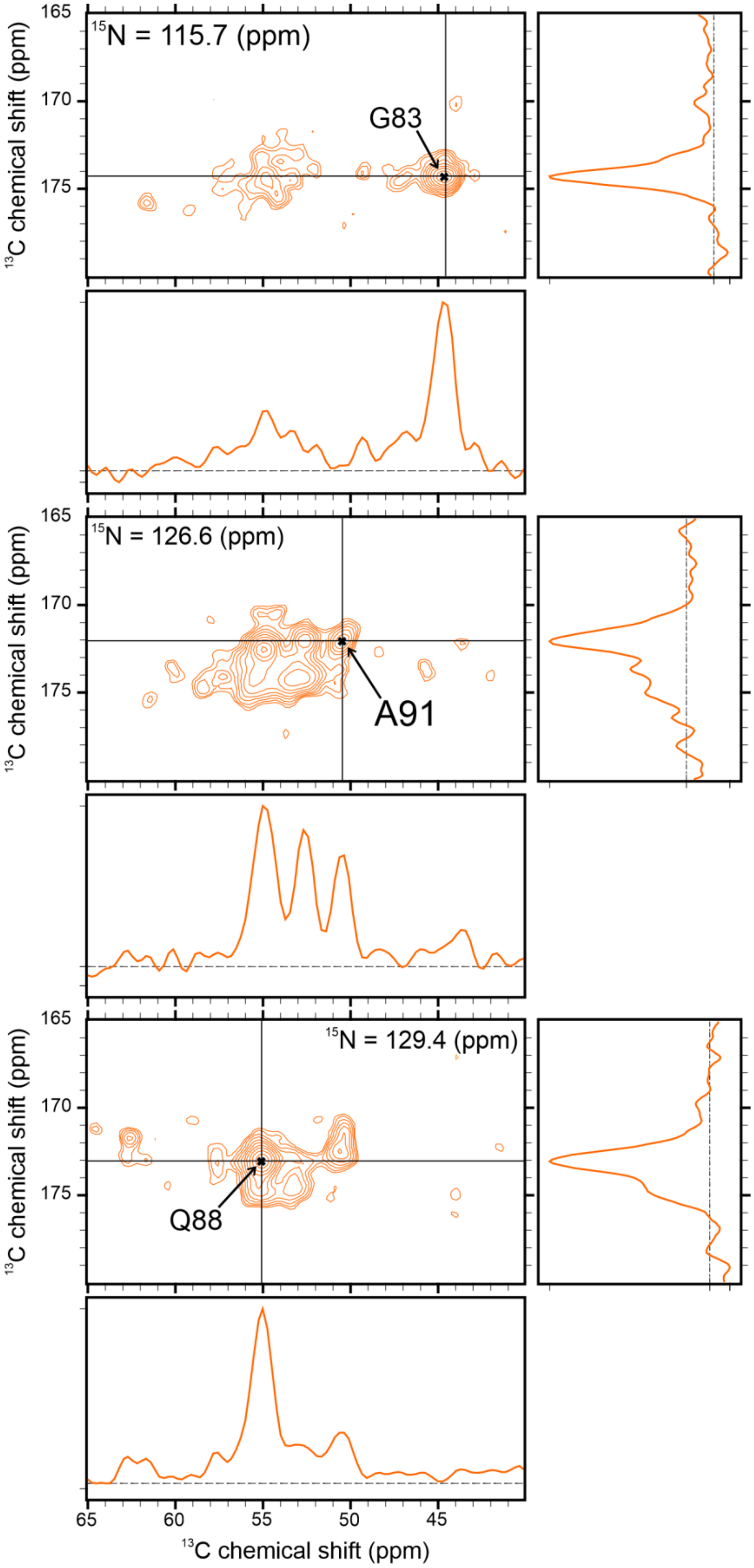

$^{15}$N = 115.7 (ppm)
G83
$^{15}$N = 126.6 (ppm)
A91
$^{15}$N = 129.4 (ppm)
Q88
$^{13}$C chemical shift (ppm)
165
170
175
65
60
55
50
45

**Supplemental Figure 4: 2D Planes from the 3D CANCO Spectrum of the Wild-type Seeded TIA1 LC Domain Fibrils Related to Figure 2.** These 2D planes demonstrate the signal to noise and increased resolution in the 3D cross polarization-based CANCO experiment. The side and bottom panels next to each plane contain slices from the data at the location of the horizontal and vertical lines. The contours are drawn at intensity values increasing by a factor of 1.3.

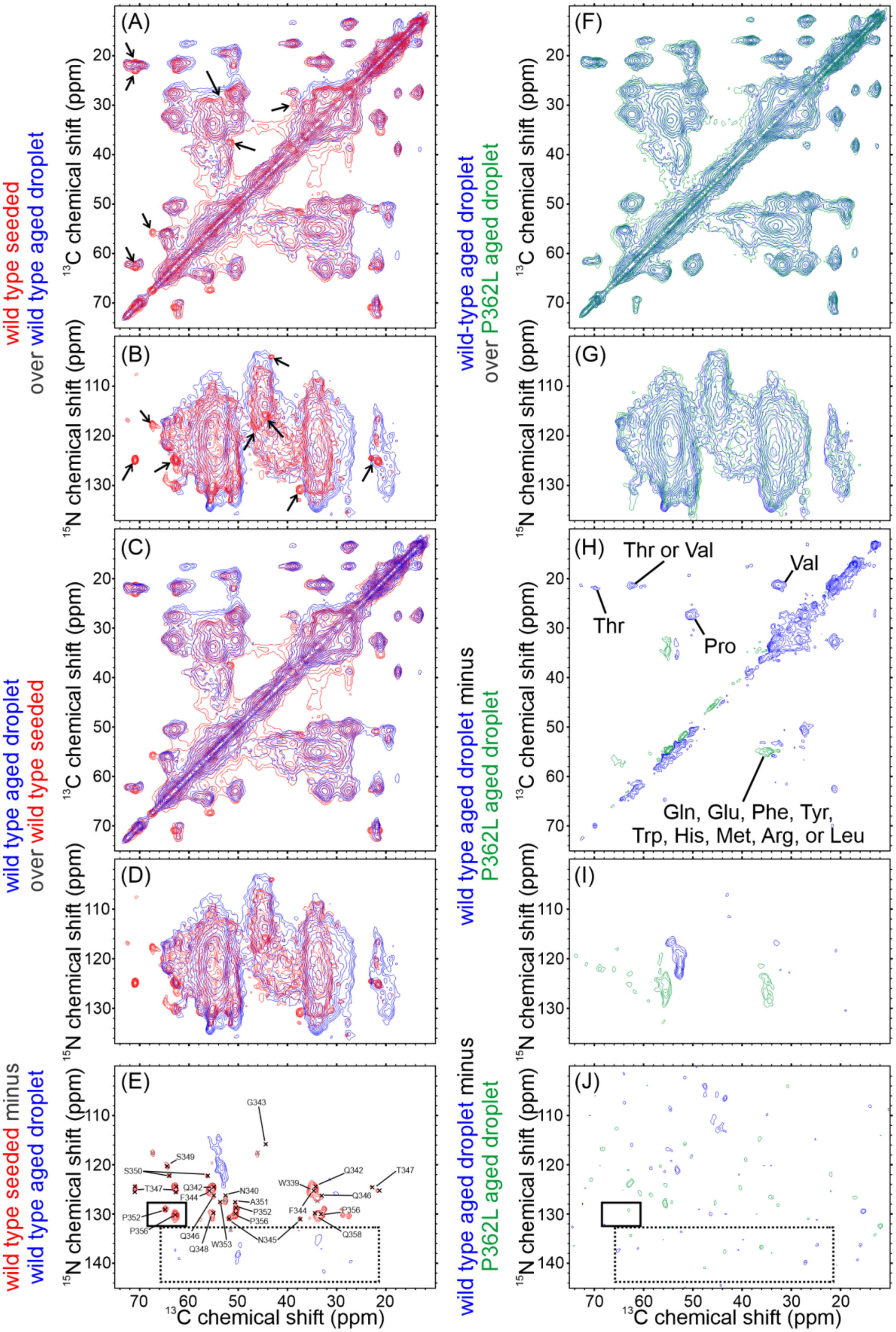
(A)
(B)
(C)
(D)
(E)
(F)
(G)
(H)
(I)
(J)
wild type seeded over wild type aged droplet
wild type aged droplet over wild type seeded
wild type seeded minus wild type aged droplet
wild-type aged droplet over P362L aged droplet
wild type aged droplet minus P362L aged droplet
wild type aged droplet minus P362L aged droplet
13C chemical shift (ppm)
15N chemical shift (ppm)
Thr or Val
Val
Thr
Pro
Gln, Glu, Phe, Tyr,
Trp, His, Met, Arg, or Leu
G343
S349
S350
T347
Q342
F344
N340
A351
P352
P356
P352
P356
Q346
Q348
W353
N345
W339
Q342
Q346
T347
F344
P356
Q358

**Supplemental Figure 5: Comparison of Seeded Fibril and Aged Liquid Droplet Spectra for Wild-type and P362L Mutant TIA1 LC Domains Related to Figure 4.** The label to the left of each spectrum indicates whether it is an overlay or difference spectrum and what two spectra were used to produce it, color coded to match the contours in the spectra. (A)–(D) overlays of the wild-type seeded and wild-type aged liquid droplet TIA1 LC domain 2D cross polarization-based spectra. Black arrows point to the locations of the most significant differences in (A) and (B). (E) difference spectrum for the TEDOR-based spectra of the wild-type seeded fibril and wild-type aged liquid droplet TIA1 LC domain samples. In (E) the blue contours represent signals that are more prominent in the aged liquid droplet sample, the red contours represent signals that are more prominent in the seeded fibril sample, the black solid box highlights the signals arising from conformationally homogenous and rigid Pro residues, and the black dashed box highlights the locations expected for the signals arising from the conformationally heterogenous but rigid Pro residues. (F) and (G) overlays of the aged liquid droplet cross polarization-based spectra from the wild-type and P362L mutant TIA1 LC domains. (H) and (I) the difference spectra calculated from the spectra in (F) and (G). The signals labeled in (H) were identified based on the statistical distributions of amino acid NMR chemical shifts in proteins. (J) the difference spectrum for the TEDOR-based spectra of the wild-type and P362L mutant aged liquid droplet TIA1 LC domain samples. In (J) the green contours represent signals that are more prominent in the P362L mutant sample, the blue contours represent signals that are more prominent in the wild-type sample, the black solid box highlights the signals arising from conformationally homogenous and rigid Pro residues, and the black dashed box highlights the locations expected for the signals arising from the conformationally heterogenous but rigid Pro residues. The contours in the carbon-carbon spectra are drawn at intensities increasing by a factor of 1.4. The contours in the nitrogen-carbon spectra are drawn at intensities increasing by a factor of 1.25.

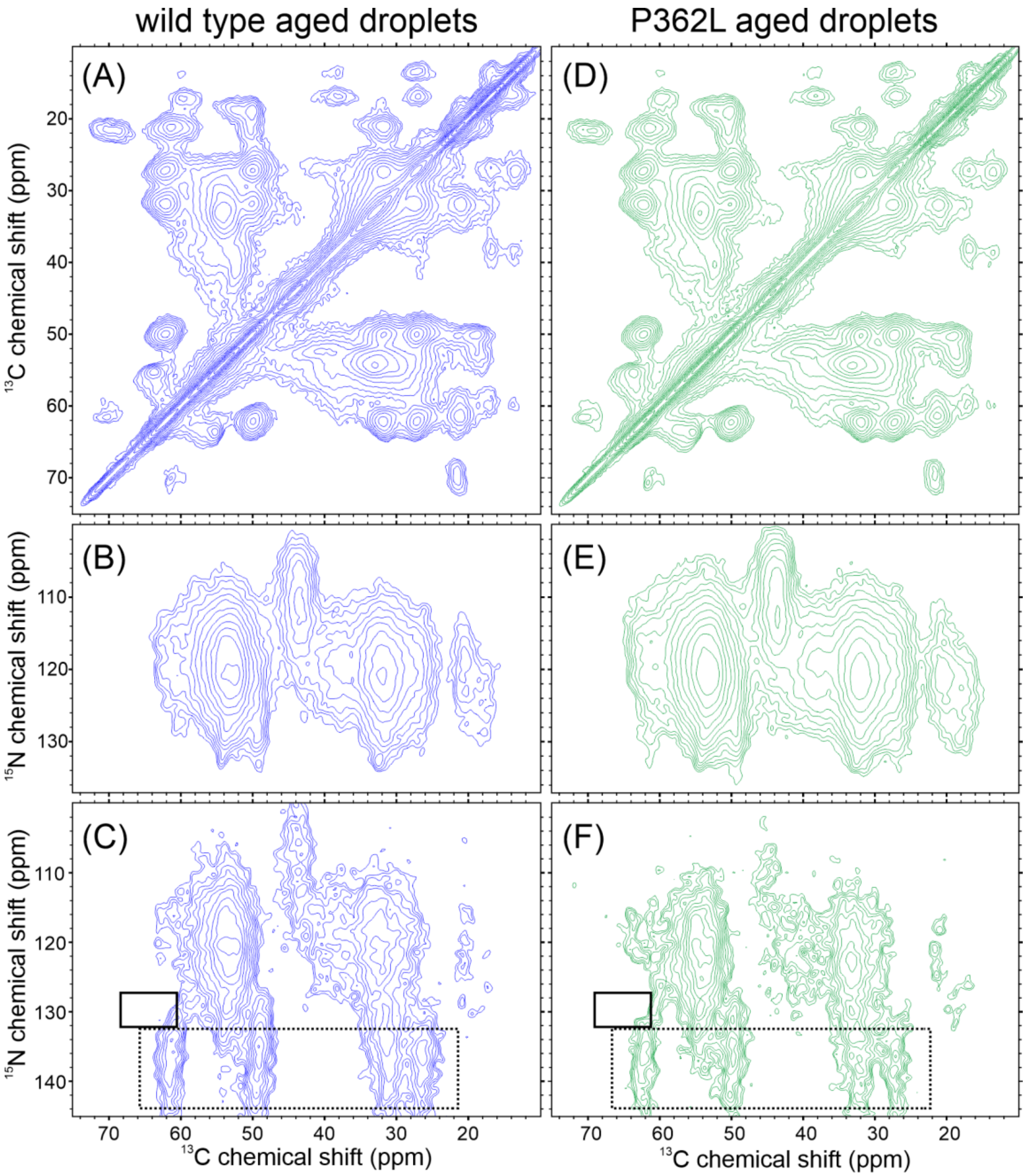


**Supplemental Figure 6: Low Temperature Spectra of the Wild-type and P362L Mutant TIA1 LC Domain Aged Droplets Related to Figure 4.** The blue contours are the wild-type TIA1 LC domain aged liquid droplets. The green contours are the P362L mutant TIA1 LC domain aged droplets. (A) and (D) are cross polarization-based carbon-carbon spectra with contours drawn at intensity levels increasing by a factor of 1.4. (B) and (E) are cross polarization-based nitrogen-carbon spectra with contours increasing by a factor of 1.25. (C) and (F) are TEDOR-based nitrogen-carbon spectra with contours increasing by a factor of 1.25. Dashed-line boxes are the signals arising from poorly ordered Pro residues. Solid-line boxes mark the locations expected for the well-ordered Pro residues present in the spectrum of the TIA1 LC domain wild-type seeded fibrils. The sample temperatures are approximately −20 °C.

**Supplemental Table S1. Solid state NMR data collection and processing parameters.**

| | Spectrum | Acquisition Parameters[a,b] | Processing Parameters[c] |
|---|---|---|---|
| **Seeded Fibrils** | 2D $^{13}C$-$^{13}C$ CP-DARR | ns=80; $\tau_{aq}$=15.36 ms; $\nu_{1H\text{-}carr}$=3.4 ppm; $\nu_{13C\text{-}carr}$=98.0 ppm; $\nu_{1H\text{-}CP}$=63 kHz; $\nu_{13C\text{-}CP}$=50 kHz; $\nu_{1Hdec}$=83.3 kHz; $\nu_{DARR}$=13 kHz; $\nu_{MAS}$=12.75 kHz; $\tau_{1H\text{-}\pi/2}$=3 µs; $\tau_{13C\text{-}\pi/2}$=4 µs; $\tau_{CP}$=1.0 ms; $\tau_{DARR}$=50 ms; $\tau_{1H\text{-}dec}$=4.9 µs; $\Delta t_1$=23.4 µs; $\tau_{t1}$=5.5 ms; $\tau_{rec}$=1.5 s; | $GLB_{t1}$=125 Hz; $GLB_{t2}$=125 Hz; LP in $t_2$ |
| | 2D CP-NCACX | ns=384; $\tau_{aq}$=15.36 ms; $\nu_{1H\text{-}carr}$=3.4 ppm; $\nu_{13C\text{-}carr}$=55.0 ppm; $\nu_{15N\text{-}carr}$=118.0 ppm; $\nu_{1H\text{-}CP}$=46 kHz; $\nu_{15N\text{-}CP}$=33 kHz; $\nu_{15N\text{-}SCP}$=36 kHz; $\nu_{13CA\text{-}SCP}$=23 kHz; $\nu_{1H\text{-}dec}$=83.3 kHz; $\nu_{DARR}$=13 kHz; $\nu_{MAS}$=12.75 kHz; $\tau_{1H\text{-}\pi/2}$=3 µs; $\tau_{13C\text{-}\pi/2}$=4 µs; $\tau_{15N\text{-}\pi/2}$=6 µs; $\tau_{CP}$=0.5 ms; $\tau_{SCP}$=5.0 ms; $\tau_{DARR}$=50 ms; $\tau_{1H\text{-}dec}$=4.9 µs; $\Delta t_1$=156.8 µs; $\tau_{t1}$=11.6 ms; $\tau_{rec}$=1.5 s; | $GLB_{t1}$=60 Hz; $GLB_{t2}$=125 Hz; |
| | 2D CP-NCOCX | ns=288; $\tau_{aq}$=15.36 ms; $\nu_{1H\text{-}carr}$=3.4 ppm; $\nu_{13C\text{-}carr}$=176.8 ppm; $\nu_{15N\text{-}carr}$=118.0 ppm; $\nu_{1H\text{-}CP}$=46 kHz; $\nu_{15N\text{-}CP}$=33 kHz; $\nu_{15N\text{-}SCP}$=36 kHz; $\nu_{13CO\text{-}SCP}$=48 kHz; $\nu_{1H\text{-}dec}$=83.3 kHz; $\nu_{DARR}$=13 kHz; $\nu_{MAS}$=12.75 kHz; $\tau_{1H\text{-}\pi/2}$=3 µs; $\tau_{13C\text{-}\pi/2}$=4 µs; $\tau_{15N\text{-}\pi/2}$=6 µs; $\tau_{CP}$=0.5 ms; $\tau_{SCP}$=5.0 ms; $\tau_{DARR}$=50 ms; $\tau_{1H\text{-}dec}$=4.9 µs; $\Delta t_1$=156.8 µs; $\tau_{t1}$=11.6 ms; $\tau_{rec}$=1.5 s; | $GLB_{t1}$=60 Hz; $GLB_{t2}$=125 Hz; |
| | 2D CP-TEDOR-NCACX | ns=800; $\tau_{aq}$=15.36 ms; $\nu_{1H\text{-}carr}$=5.2 ppm; $\nu_{13C\text{-}carr}$=57.0 ppm; $\nu_{15N\text{-}carr}$=120.0 ppm; $\nu_{1H\text{-}CP}$=63 kHz; $\nu_{13C\text{-}CP}$=50 kHz; $\nu_{DARR}$=13 kHz; $\nu_{1Hdec}$=83.3 kHz; $\nu_{MAS}$=12.7 kHz; $\tau_{1H\text{-}\pi/2}$=3 µs; $\tau_{13C\text{-}\pi/2}$=4 µs; $\tau_{13C\text{-}soft}$=431 µs; $\tau_{15N\text{-}\pi/2}$=6 µs; $\tau_{TEDOR}$=1.6 ms; $\tau_{z\text{-}filter}$=78.4 µs; $\tau_{DARR}$=50 ms; $\tau_{1H\text{-}dec}$=6.3 µs; $\Delta t_1$=156.9 µs; $\tau_{t1}$=11.6 ms; $\tau_{rec}$=1.5 s; | $GLB_{t1}$=60 Hz; $GLB_{t2}$=125 Hz; |
| | 2D $^{1}H$-$^{13}C$ INEPT | ns=16; $\tau_{aq}$=20.48 ms; $\nu_{1H\text{-}carr}$=5.2 ppm; $\nu_{13C\text{-}carr}$=67.3 ppm; $\nu_{1H\text{-}dec}$=25.0 kHz; $\nu_{MAS}$=12.75 kHz; $\tau_{1H\text{-}\pi/2}$=3 µs; $\tau_{13C\text{-}\pi/2}$=4 µs; $\tau_{J/2}$=1.2 ms; $\Delta t_1$=75.0 µs; $\tau_{t1}$=11.25 ms; $\tau_{rec}$=1.5 s; | $GLB_{t1}$=10 Hz; $GLB_{t2}$=50 Hz; |
| | 3D CP-NCACX | ns=16; $\tau_{aq}$=10.24 ms; $\nu_{1H\text{-}carr}$=2.3 ppm; $\nu_{13C\text{-}carr}$=53.4 ppm; $\nu_{15N\text{-}carr}$=121.6 ppm; $\nu_{1H\text{-}CP}$=50 kHz; $\nu_{15N\text{-}CP}$=37 kHz; $\nu_{15N\text{-}SCP}$=33 kHz; $\nu_{13CA\text{-}SCP}$=20 kHz; $\nu_{1H\text{-}dec}$=83.3 kHz; $\nu_{DARR}$=13 kHz; $\nu_{MAS}$=12.65 kHz; $\tau_{1H\text{-}\pi/2}$=3 µs; $\tau_{13C\text{-}\pi/2}$=4 µs; $\tau_{15N\text{-}\pi/2}$=6 µs; $\tau_{CP}$=0.5 ms; $\tau_{SCP}$=5.0 ms; $\tau_{DARR}$=50 ms; $\tau_{1H\text{-}dec}$=6.3 µs; $\Delta t_2$=88.2 µs; $\tau_{t2}$=4.4 ms; $\Delta t_1$=138.6 µs; $\tau_{t1}$=10.1 ms; $\tau_{rec}$=1.5 s; | $GLB_{t1}$=60 Hz; $GLB_{t2}$=125 Hz; $GLB_{t3}$=125 Hz; |

| | | | |
|---|---|---|---|
| | 3D CP-NCOCX | ns=32; $\tau_{aq}$=10.24 ms; $\nu_{1H-carr}$=2.3 ppm; $\nu_{13C-carr}$=175.4 ppm; $\nu_{15N-carr}$=121.6 ppm; $\nu_{1H-CP}$=50 kHz; $\nu_{15N-CP}$=37 kHz; $\nu_{15N-SCP}$=33 kHz; $\nu_{13CO-SCP}$=46 kHz; $\nu_{1H-dec}$=83.3 kHz; $\nu_{DARR}$=13 kHz; $\nu_{MAS}$=12.65 kHz; $\tau_{1H-\pi/2}$=3 µs; $\tau_{13C-\pi/2}$=4 µs; $\tau_{15N-\pi/2}$=6 µs; $\tau_{CP}$=0.5 ms; $\tau_{SCP}$=5.0 ms; $\tau_{DARR}$=50 ms; $\tau_{1H-dec}$=6.3 µs; $\Delta t_2$=138.6 µs; $\tau_{t2}$=4.4 ms; $\Delta t_1$=138.6 µs; $\tau_{t1}$=10.1 ms; $\tau_{rec}$=1.5 s; | $GLB_{t1}$=60 Hz; $GLB_{t2}$=125 Hz; $GLB_{t3}$=125 Hz; |
| | 3D CP-CANCO | ns=28; $\tau_{aq}$=15.36 ms; $\nu_{1H-carr}$=1.0 ppm; $\nu_{13C-carr}$=55.0 ppm; $\nu_{15N-carr}$=118.0 ppm; $\nu_{1H-CP}$=59 kHz; $\nu_{13C-CP}$=46 kHz; $\nu_{15N-SCP}$=34 kHz; $\nu_{13CA-SCP}$=22 kHz; $\nu_{13CO-SCP-eff}$=47 kHz; $\nu_{1Hdec}$=83.3 kHz; $\nu_{MAS}$=12.7 kHz; $\tau_{1H-\pi/2}$=3 µs; $\tau_{SCP}$=4 ms; $\tau_{1H-dec}$=6 µs; $\Delta t_2$=151.2 µs; $\tau_{t2}$=8.1 ms; $\Delta t_1$=75.6 µs; $\tau_{t1}$=4.2 ms; $\tau_{rec}$=1.5 s; | $GLB_{t1}$=125 Hz; $GLB_{t2}$=60 Hz; $GLB_{t3}$=125 Hz; |
| **Aged Droplet (WT and P36L Mutant)** | 2D $^{13}C$-$^{13}C$ CP-DARR | $ns_{WT,263K}$=80; $ns_{WT,233K}$=96; $ns_{P362L,263K}$=64; $ns_{P362L,233K}$=80; $\tau_{aq}$=7.68 ms; $\nu_{1H-carr}$=3.9 ppm; $\nu_{13C-carr}$=99.2 ppm; $\nu_{1H-CP}$=63 kHz; $\nu_{13C-CP}$=50 kHz; $\nu_{1Hdec}$=83.3 kHz; $\nu_{DARR}$=13 kHz; $\nu_{MAS}$=12.75 kHz; $\tau_{1H-\pi/2}$=3 µs; $\tau_{13C-\pi/2}$=4 µs; $\tau_{CP}$=1.0 ms; $\tau_{DARR}$=50 ms; $\tau_{1H-dec}$=6.0 µs; $\Delta t_1$=12.0 µs; $\tau_{t1,263K}$=3.9 ms; $\tau_{t1,233K}$=3.5 ms; $\tau_{rec}$=2.0 s; | $GLB_{t1}$=75 Hz; $GLB_{t2}$=75 Hz; LP in $t_2$ |
| | 2D CP-NCACX | $ns_{WT,263K}$=616; $ns_{WT,233K}$=288; $ns_{P362L,263K}$=256; $ns_{P362L,233K}$=288; $\tau_{aq}$=10.24 ms; $\nu_{1H-carr}$=3.9 ppm; $\nu_{13C-carr}$=63.8 ppm; $\nu_{15N-carr}$=120.4 ppm; $\nu_{1H-CP}$=61 kHz; $\nu_{15N-CP}$=49 kHz; $\nu_{15N-SCP}$=34 kHz; $\nu_{13CA-SCP}$=22 kHz; $\nu_{1H-dec}$=83.3 kHz; $\nu_{DARR}$=13 kHz; $\nu_{MAS}$=12.75 kHz;$\tau_{1H-\pi/2}$=3 µs; $\tau_{13C-\pi/2}$=4 µs; $\tau_{15N-\pi/2}$=6 µs; $\tau_{CP}$=0.75 ms; $\tau_{SCP}$=4.0 ms; $\tau_{DARR}$=50 ms; $\tau_{1H-dec}$=6.0 µs; $\Delta t_1$=108.0 µs; $\tau_{t1,263K}$=10.0 ms; $\tau_{t1,233K}$=10.0 ms; $\tau_{rec}$=2.0 s; | $GLB_{t1}$=80 Hz; $GLB_{t2}$=150 Hz; |
| | 2D CP-TEDOR-NCACX | $ns_{WT,233K}$=512; $ns_{P362L,233K}$=576; $\tau_{aq}$=7.68 ms; $\nu_{1H-carr}$=3.9 ppm; $\nu_{13C-carr}$=58.1 ppm; $\nu_{15N-carr}$=120.7 ppm; $\nu_{1H-CP}$=63 kHz; $\nu_{13C-CP}$=50 kHz; $\nu_{DARR}$=13 kHz; $\nu_{1Hdec}$=83.3 kHz; $\nu_{MAS}$=12.75 kHz; $\tau_{1H-\pi/2}$=3 µs; $\tau_{13C-\pi/2}$=4 µs; $\tau_{13C-soft}$=431 µs; $\tau_{15N-\pi/2}$=6 µs; $\tau_{TEDOR}$=1.6 ms; $\tau_{z-filter}$=78.4 µs; $\tau_{DARR}$=50 ms; $\tau_{1H-dec}$=6.0 µs; $\Delta t_1$=156.8 µs; $\tau_{t1}$=8.3 ms; $\tau_{rec}$=2.0 s; | $GLB_{t1}$=80 Hz; $GLB_{t2}$=150 Hz; |

[a] All $^{1}H$-X CP transfers used a 20% ramp on the $^{1}H$ channel. All $^{15}N$-$^{13}C$ Specific-CP transfers used a 5% ramp on the $^{15}N$ channel.

[b] ns, number of scans averaged (subscripts for the aged droplet samples specify the sample and temperature); $\tau_{aq}$, total acquisition time in the direct dimension; $\nu_{1H-carr}$, $^{1}H$ carrier frequency; $\nu_{13C-carr}$, $^{13}C$ carrier frequency; $\nu_{15N-carr}$, $^{15}N$ carrier frequency; $\nu_{1H-CP}$, $^{1}H$ CP pulse power; $\nu_{13C-CP}$, $^{13}C$ CP pulse power; $\nu_{15N-CP}$, $^{15}N$ CP pulse power; $\nu_{15N-SCP}$, $^{15}N$ Specific-CP pulse power; $\nu_{13CA-}$

$_{SCP}$, $^{13}$C α-carbon Specific-CP pulse power; $\nu_{13CO\text{-}SCP}$, $^{13}$C carbonyl-carbon Specific-CP pulse power; $\nu_{13CO\text{-}SCP\text{-}eff}$, $^{13}$C carbonyl-carbon Specific-CP effective field pulse power; $\nu_{1H\text{-}dec}$, $^{1}$H decoupling pulse power; $\nu_{DARR}$, $^{13}$C-$^{13}$C DARR $^{1}$H pulse power; $\nu_{MAS}$, magic angle spinning frequency; $\tau_{1H\text{-}\pi/2}$, $^{1}$H π/2 pulse length; $\tau_{13C\text{-}\pi/2}$, $^{13}$C π/2 pulse length; $\tau_{13C\text{-}soft}$, $^{13}$C selective pulse length; $\tau_{15N\text{-}\pi/2}$, $^{15}$N π/2 pulse length; $\tau_{CP}$, $^{1}$H-X CP contact time; $\tau_{SCP}$, $^{15}$N-$^{13}$C Specific-CP contact time; $\tau_{DARR}$, $^{13}$C-$^{13}$C DARR mixing time; $\tau_{J/2}$,1/2 echo delay; $\tau_{1H\text{-}dec}$, $^{1}$H SPINAL64 π pulse length; $\Delta t_2$, time increment in the $t_2$ dimension; $\tau_{t2}$, total acquisition time in the $t_2$ dimension; $\Delta t_1$, time increment in the $t_1$ dimension; $\tau_{t1}$, total acquisition time in the $t_1$ dimension (subscripts for the aged droplet samples specify the temperature); $\tau_{rec}$, recycle delay; $\tau_{z\text{-}filter}$, z-filter time for the TEDOR experiments; $\tau_{TEDOR}$ total TEDOR mixing time;

[c] Zero filling was applied twice in each dimension prior to Fourier transformation. GLB, gaussian line broadening and LP, forward linear prediction with standard NMRPipe parameters.

**Supplemental Table S2. Assigned solid state NMR chemical shifts (ppm).** Signals in the table represent the average from the NCACX, NCOCX, and CANCO 3D spectra.

| **Unambiguously Assigned Chemical Shifts** | | | | | | |
|---|---|---|---|---|---|---|
| Residue | N | CA | C | CB | CG | CD |
| Q337 | | | 174.0 | | | |
| A338 | 120.7 | 51.0 | 175.0 | 21.9 | | |
| W339 | 125.4 | 54.0 | 174.3 | 34.2 | | |
| N340 | 126.3 | 52.6 | 173.7 | 41.5 | | |
| Q341 | 119.8 | 55.1 | 175.9 | 35.4 | 35.4 | 179.2 |
| Q342 | 124.4 | 54.9 | 174.3 | 32.0 | 34.5 | 177.0 |
| G343 | 116.0 | 44.5 | 169.1 | | | |
| F344 | 125.5 | 55.8 | 174.6 | 34.9 | | |
| N345 | 131.0 | 51.9 | 172.5 | 37.5 | 174.9 | |
| Q346 | 126.5 | 55.0 | 174.7 | 33.3 | 33.2 | 178.4 |
| T347 | 125.1 | 62.5 | 173.5 | 71.0 | 21.5/23.0[a] | |
| Q348 | 130.0 | 55.0 | 173.3 | 33.2 | 34.8 | |
| S349 | 120.6 | 55.8 | 174.8 | 64.3 | | |
| S350 | 122.2 | 56.2 | 172.1 | 63.9 | | |
| A351 | 127.1 | 50.6 | 171.2 | 18.6 | | |
| P352 | 129.0 | 64.7 | 174.3 | 32.5 | 29.1 | 50.3 |
| W353 | 127.5 | 53.6 | 174.3 | 32.0 | 111.5 | |
| M354 | 125.3 | 53.2 | 173.5 | 38.4 | 29.7 | |
| G355 | 104.1 | 43.3 | 171.8 | | | |
| P356 | 130.1 | 62.7 | 173.4 | 32.9 | 28.6 | 50.5 |
| N357 | 118.5 | 52.4 | 173.5 | 41.3 | 175.2 | |
| **XGXX Motif (ppm)** | | | | | | |
| X | | | 176.1 | | | |
| G | 117.8 | 46.0 | 169.9 | | | |
| X | 122.9 | 54.0 | 171.9 | 35.8 | | |
| X | 127.3 | 52.6 | 174.0 | 34.1 | | |

[a]Two Thr signals are present with nearly identical CA and CB chemical shifts but differing CG2 chemical shifts.

**Supplemental Table S3. NCACX MCASSIGN Table.** Chemical shifts and uncertainties are in ppm. Amino acid types are single letter codes. A value of '1111.1' is used to indicate no signal.

| N | CA | CO | CB | CG | CD | ΔN | ΔCA | ΔCO | ΔCB | ΔCG | ΔCD | Amino Acid Type |
|---|---|---|---|---|---|---|---|---|---|---|---|---|
| 104.2 | 43.2 | 171.8 | 1111.1 | 1111.1 | 1111.1 | 0.4 | 0.3 | 0.3 | 0.3 | 0.3 | 0.3 | G |
| 113.8 | 53.0 | 174 | 44.1 | 177.1 | 1111.1 | 0.4 | 0.4 | 0.4 | 0.4 | 0.4 | 0.3 | ND |
| 116.0 | 44.4 | 169.2 | 1111.1 | 1111.1 | 1111.1 | 0.4 | 0.4 | 0.6 | 0.3 | 0.3 | 0.3 | G |
| 117.7 | 45.9 | 169.8 | 1111.1 | 1111.1 | 1111.1 | 0.4 | 0.4 | 0.6 | 0.3 | 0.3 | 0.3 | G |
| 117.7 | 55.7 | 174.2 | 67.2 | 1111.1 | 1111.1 | 0.4 | 0.3 | 0.8 | 0.3 | 0.3 | 0.3 | S |
| 117.8 | 55.8 | 174.3 | 64.7 | 1111.1 | 1111.1 | 0.4 | 0.3 | 0.8 | 0.3 | 0.3 | 0.3 | S |
| 118.6 | 52.4 | 173.7 | 41.4 | 175.2 | 1111.1 | 0.4 | 0.3 | 0.5 | 0.3 | 0.6 | 0.3 | ND |
| 119.3 | 55.2 | 175 | 34.3 | 1111.1 | 1111.1 | 0.5 | 0.3 | 1.5 | 0.5 | 1.2 | 0.5 | RDNQEHILMFWY |
| 119.6 | 55.1 | 175.8 | 35.5 | 1111.1 | 1111.1 | 0.4 | 0.3 | 1.5 | 0.4 | 0.3 | 0.3 | RDNQEHILMFWY |
| 120.6 | 55.8 | 174.2 | 64.3 | 1111.1 | 1111.1 | 0.4 | 0.3 | 1.1 | 0.3 | 0.3 | 0.3 | S |
| 120.7 | 50.9 | 175 | 21.8 | 1111.1 | 1111.1 | 0.4 | 0.3 | 0.5 | 0.3 | 0.3 | 0.3 | A |
| 120.8 | 55.4 | 174.5 | 33.8 | 1111.1 | 1111.1 | 0.5 | 0.4 | 1.5 | 0.8 | 0.3 | 0.3 | RDNQEHILMFWY |
| 122.3 | 56.2 | 172.3 | 63.9 | 1111.1 | 1111.1 | 0.4 | 0.3 | 0.3 | 0.3 | 0.3 | 0.3 | S |
| 122.4 | 55.3 | 174.4 | 33.9 | 1111.1 | 1111.1 | 0.4 | 0.5 | 0.5 | 1.5 | 0.3 | 0.3 | RDNQEHILMFWY |
| 122.9 | 54.0 | 171.8 | 35.7 | 1111.1 | 1111.1 | 0.4 | 0.3 | 0.3 | 0.7 | 0.3 | 0.3 | RDNQEHILMFWY |
| 124.5 | 54.9 | 174.3 | 31.9 | 34.3 | 176.9 | 0.5 | 0.4 | 0.4 | 0.7 | 0.7 | 0.7 | QE |
| 124.5 | 62.6 | 172.8 | 70.9 | 23.0 | 1111.1 | 0.4 | 0.3 | 0.3 | 0.3 | 0.3 | 0.3 | T |
| 124.5 | 54.9 | 174.2 | 34.5 | 1111.1 | 1111.1 | 0.5 | 0.4 | 0.8 | 0.7 | 0.7 | 0.8 | RDNQEHILMFWY |
| 125.1 | 53.1 | 173.6 | 38.5 | 29.5 | 1111.1 | 0.4 | 0.3 | 1.0 | 0.3 | 0.3 | 0.3 | RDNQEHILMFWY |
| 125.2 | 55.8 | 174.3 | 34.7 | 1111.1 | 1111.1 | 0.5 | 0.4 | 0.7 | 0.7 | 0.3 | 0.3 | RDNQEHILMFWY |
| 125.3 | 62.6 | 172.8 | 70.9 | 21.4 | 1111.1 | 0.4 | 0.3 | 0.3 | 0.3 | 0.3 | 0.3 | T |
| 125.4 | 54.0 | 173.1 | 34.8 | 1111.1 | 1111.1 | 0.6 | 0.4 | 1.2 | 1.0 | 0.3 | 0.3 | RDNQEHILMFWY |
| 126.2 | 52.4 | 173.6 | 41.4 | 1111.1 | 1111.1 | 0.4 | 0.3 | 0.4 | 0.3 | 0.3 | 0.3 | RDNQEHILMFWY |
| 126.5 | 55.0 | 174.7 | 33.2 | 33.2 | 178.4 | 0.4 | 0.3 | 0.3 | 1.6 | 1.6 | 0.3 | QE |
| 127.3 | 52.6 | 174 | 34.1 | 1111.1 | 1111.1 | 0.4 | 0.3 | 0.4 | 0.3 | 0.3 | 0.3 | RDNQEHILMFWY |
| 127.4 | 50.5 | 171.2 | 18.5 | 1111.1 | 1111.1 | 0.4 | 0.3 | 0.3 | 0.3 | 0.3 | 0.3 | A |
| 127.5 | 53.7 | 174.1 | 31.9 | 1111.1 | 1111.1 | 0.4 | 0.3 | 0.7 | 0.3 | 0.3 | 0.3 | RDNQEHILMFWY |
| 129.1 | 64.7 | 174.2 | 32.5 | 29.0 | 50.2 | 0.4 | 0.3 | 0.7 | 0.3 | 0.3 | 0.3 | P |
| 129.8 | 55.0 | 173.2 | 32.7 | 34.5 | 1111.1 | 0.4 | 0.3 | 0.4 | 0.7 | 0.7 | 0.3 | RDNQEHILMFWY |
| 130.3 | 62.8 | 173.7 | 32.9 | 28.2 | 50.6 | 0.4 | 0.3 | 1.7 | 1.4 | 0.9 | 0.4 | P |
| 130.9 | 51.8 | 172.4 | 37.4 | 175.0 | 1111.1 | 0.4 | 0.3 | 0.3 | 0.3 | 0.3 | 0.3 | ND |
| 133.0 | 55.5 | 173.9 | 34.0 | 1111.1 | 1111.1 | 0.5 | 0.4 | 0.4 | 0.7 | 0.3 | 0.3 | RDNQEHILMFWY |
| 133.2 | 51.4 | 172.5 | 37.6 | 175.8 | 1111.1 | 0.4 | 0.3 | 0.3 | 0.3 | 0.3 | 0.3 | ND |

**Supplemental Table S4. NCOCX MCASSIGN Table.** Chemical shifts and uncertainties are in ppm. Amino acid types are single letter codes. A value of '1111.1' is used to indicate no signal.

| N | CA | CO | CB | CG | CD | ΔN | ΔCA | ΔCO | ΔCB | ΔCG | ΔCD | Amino Acid Type |
|---|---|---|---|---|---|---|---|---|---|---|---|---|
| 104.0 | 53.2 | 173.4 | 38.5 | 29.7 | 1111 | 0.3 | 0.3 | 0.3 | 0.3 | 0.3 | 0.4 | RDNQEHILMFWY |
| 113.6 | 43.9 | 171.7 | 1111.1 | 1111.1 | 1111.1 | 0.3 | 0.3 | 0.3 | 0.4 | 0.4 | 0.4 | G |
| 116.0 | 55.0 | 174.3 | 32.0 | 34.4 | 177.1 | 0.3 | 0.3 | 0.3 | 0.5 | 0.3 | 0.4 | QE |
| 117.5 | 55.3 | 174.6 | 33.1 | 1111.1 | 1111.1 | 0.4 | 0.3 | 0.3 | 0.3 | 0.4 | 0.4 | RDNQEHILMFWY |
| 118.3 | 62.8 | 173.1 | 32.9 | 29.0 | 50.4 | 0.4 | 0.4 | 0.4 | 0.4 | 0.4 | 0.4 | P |
| 119.5 | 55.6 | 174.7 | 34.1 | 1111.1 | 1111.1 | 0.5 | 0.4 | 0.4 | 2.0 | 0.4 | 0.4 | RDNQEHILMFWY |
| 119.7 | 52.5 | 173.6 | 41.5 | 1111.1 | 1111.1 | 0.4 | 0.3 | 0.3 | 0.3 | 0.4 | 0.4 | RDNQEHILMFWY |
| 120.7 | 55.0 | 173.3 | 33.9 | 1111.1 | 1111.1 | 0.4 | 0.4 | 0.3 | 1.5 | 0.4 | 0.4 | RDNQEHILMFWY |
| 121.0 | 52.7 | 173.6 | 33.8 | 1111.1 | 1111.1 | 0.4 | 0.3 | 0.3 | 1.5 | 0.4 | 0.4 | RDNQEHILMFWY |
| 122.2 | 44.4 | 172.4 | 1111.1 | 1111.1 | 1111.1 | 0.4 | 0.7 | 0.7 | 0.4 | 0.4 | 0.4 | G |
| 122.3 | 55.7 | 174.8 | 64.2 | 1111.1 | 1111.1 | 0.3 | 0.3 | 0.3 | 0.3 | 0.4 | 0.4 | S |
| 122.3 | 55.0 | 174 | 33.7 | 1111.1 | 1111.1 | 0.5 | 0.7 | 1.0 | 2.0 | 0.4 | 0.4 | RDNQEHILMFWY |
| 122.9 | 46.0 | 169.9 | 1111.1 | 1111.1 | 1111.1 | 0.3 | 0.6 | 0.6 | 0.4 | 0.4 | 0.4 | G |
| 123.5 | 52.8 | 172.6 | 1111.1 | 1111.1 | 1111.1 | 0.4 | 0.8 | 0.4 | 0.4 | 0.4 | 0.4 | RDNQEHILMFWYS |
| 123.6 | 53.7 | 173.8 | 33.6 | 1111.1 | 1111.1 | 0.4 | 0.4 | 0.4 | 2.0 | 0.4 | 0.4 | RDNQEHILMFWY |
| 123.7 | 55.3 | 174.1 | 33.6 | 1111.1 | 1111.1 | 0.4 | 0.4 | 0.4 | 2.0 | 0.4 | 0.4 | RDNQEHILMFWY |
| 124.3 | 55.1 | 176.3 | 35.1 | 1111.1 | 1111.1 | 0.4 | 0.3 | 0.4 | 0.5 | 0.4 | 0.4 | RDNQEHILMFWY |
| 124.9 | 55.1 | 174.7 | 33.3 | 1111.1 | 1111.1 | 0.4 | 0.4 | 0.4 | 1.5 | 0.4 | 0.4 | RDNQEHILMFWY |
| 125.2 | 53.6 | 174.3 | 32.5 | 1111.1 | 1111.1 | 0.4 | 0.4 | 0.4 | 1.0 | 0.4 | 0.4 | RDNQEHILMFWY |
| 125.4 | 51.0 | 175.1 | 21.8 | 1111.1 | 1111.1 | 0.4 | 0.3 | 0.3 | 0.3 | 0.4 | 0.4 | A |
| 125.6 | 44.5 | 169.2 | 1111.1 | 1111.1 | 1111.1 | 0.3 | 0.3 | 0.5 | 0.4 | 0.4 | 0.4 | G |
| 126.1 | 54.9 | 172.6 | 33.4 | 1111.1 | 1111.1 | 0.4 | 0.4 | 0.4 | 0.8 | 0.4 | 0.4 | RDNQEHILMFWY |
| 126.4 | 53.6 | 174.3 | 34.1 | 1111.1 | 1111.1 | 0.4 | 0.8 | 0.4 | 0.8 | 0.4 | 0.4 | RDNQEHILMFWY |
| 126.5 | 51.8 | 172.3 | 37.5 | 174.8 | 1111.1 | 0.3 | 0.3 | 0.3 | 0.3 | 0.3 | 0.4 | ND |
| 126.5 | 50.9 | 174.4 | 32.3 | 1111.1 | 1111.1 | 0.4 | 0.4 | 0.4 | 0.4 | 0.4 | 0.4 | RDNQEHILMFWY |
| 127.2 | 56.1 | 172.1 | 63.7 | 1111.1 | 1111.1 | 0.4 | 0.4 | 0.4 | 0.3 | 0.4 | 0.4 | S |
| 127.3 | 54.0 | 171.9 | 35.6 | 1111.1 | 1111.1 | 0.3 | 0.3 | 0.3 | 1.5 | 0.4 | 0.4 | RDNQEHILMFWY |
| 127.4 | 64.8 | 174.3 | 32.4 | 29.2 | 50.3 | 0.3 | 0.3 | 0.3 | 0.3 | 0.3 | 0.3 | P |
| 128.5 | 52.7 | 174.2 | 34.3 | 1111.1 | 1111.1 | 0.4 | 0.3 | 0.3 | 0.3 | 0.4 | 0.4 | RDNQEHILMFWY |
| 128.7 | 56.3 | 173.0 | 1111.1 | 1111.1 | 1111.1 | 0.4 | 0.3 | 0.3 | 0.4 | 0.4 | 0.4 | RDNQEHILMFWYS |
| 129.4 | 62.7 | 173.0 | 70.9 | 23.0 | 1111.1 | 0.4 | 0.3 | 0.3 | 0.3 | 0.3 | 0.3 | T |
| 129.7 | 55.7 | 174.8 | 34.4 | 1111.1 | 1111.1 | 0.4 | 0.5 | 0.4 | 1.0 | 0.4 | 0.4 | RDNQEHILMFWY |
| 129.7 | 55.3 | 173.0 | 33.9 | 1111.1 | 1111.1 | 0.4 | 0.7 | 0.4 | 0.7 | 0.4 | 0.4 | RDNQEHILMFWY |
| 130.3 | 62.5 | 172.8 | 71.0 | 21.4 | 1111.1 | 0.5 | 0.3 | 0.3 | 0.3 | 0.3 | 0.4 | T |
| 130.7 | 56.3 | 172.6 | 34.0 | 1111.1 | 1111.1 | 0.4 | 0.5 | 0.4 | 0.5 | 0.4 | 0.4 | RDNQEHILMFWY |
| 131.0 | 55.7 | 175.0 | 34.9 | 1111.1 | 1111.1 | 0.4 | 0.3 | 0.3 | 0.3 | 0.4 | 0.4 | RDNQEHILMFWY |
| 133.4 | 53.3 | 172.3 | 1111.1 | 1111.1 | 1111.1 | 0.4 | 0.4 | 0.4 | 0.4 | 0.4 | 0.4 | RDNQEHILMFWYS |

**Supplemental Table S5. CANCO MCASSIGN Table.** Chemical shifts and uncertainties are in ppm. Amino acid types are single letter codes. A value of '1111.1' is used to indicate no signal.

| N | CA | CO | CB | CG | CD | ΔN | ΔCA | ΔCO | ΔCB | ΔCG | ΔCD | Amino Acid Type |
|---|---|---|---|---|---|---|---|---|---|---|---|---|
| 104.2 | 43.3 | 173.6 | 1111.1 | 1111.1 | 1111.1 | 0.4 | 0.7 | 0.4 | 0.4 | 0.4 | 0.4 | G |
| 115.9 | 44.6 | 174.3 | 1111.1 | 1111.1 | 1111.1 | 0.4 | 0.3 | 0.3 | 0.4 | 0.4 | 0.4 | G |
| 117.5 | 55.6 | 174.6 | 1111.1 | 1111.1 | 1111.1 | 0.4 | 0.3 | 0.3 | 0.4 | 0.4 | 0.4 | S |
| 117.6 | 45.9 | 176.3 | 1111.1 | 1111.1 | 1111.1 | 0.4 | 0.3 | 0.3 | 0.4 | 0.4 | 0.4 | G |
| 118.5 | 52.5 | 173.3 | 1111.1 | 1111.1 | 1111.1 | 0.4 | 0.3 | 0.3 | 0.4 | 0.4 | 0.4 | RDNQEHILMFWY |
| 119.5 | 55.1 | 174.0 | 1111.1 | 1111.1 | 1111.1 | 0.4 | 0.4 | 2.0 | 0.4 | 0.4 | 0.4 | RDNQEHILMFWY |
| 120.5 | 55.9 | 173.3 | 1111.1 | 1111.1 | 1111.1 | 0.4 | 0.4 | 0.4 | 0.4 | 0.4 | 0.4 | S |
| 120.7 | 53.2 | 174.3 | 1111.1 | 1111.1 | 1111.1 | 0.4 | 0.3 | 0.3 | 0.4 | 0.4 | 0.4 | RDNQEHILMFWY |
| 120.8 | 51.0 | 174.1 | 1111.1 | 1111.1 | 1111.1 | 0.4 | 0.3 | 0.3 | 0.4 | 0.4 | 0.4 | A |
| 121.4 | 53.2 | 173.5 | 1111.1 | 1111.1 | 1111.1 | 0.4 | 0.4 | 0.7 | 0.4 | 0.4 | 0.4 | RDNQEHILMFWY |
| 122.0 | 54.6 | 174.6 | 1111.1 | 1111.1 | 1111.1 | 0.4 | 0.4 | 0.4 | 0.4 | 0.4 | 0.4 | RDNQEHILMFWY |
| 122.2 | 56.1 | 174.8 | 1111.1 | 1111.1 | 1111.1 | 0.4 | 0.4 | 0.4 | 0.4 | 0.4 | 0.4 | S |
| 122.3 | 55.4 | 173.6 | 1111.1 | 1111.1 | 1111.1 | 0.8 | 0.8 | 1.5 | 0.4 | 0.4 | 0.4 | RDNQEHILMFWY |
| 122.8 | 51.1 | 174.2 | 1111.1 | 1111.1 | 1111.1 | 0.4 | 0.3 | 0.3 | 0.4 | 0.4 | 0.4 | RDNQEHILMFWY |
| 122.9 | 54.0 | 170.0 | 1111.1 | 1111.1 | 1111.1 | 0.4 | 0.3 | 1.0 | 0.4 | 0.4 | 0.4 | RDNQEHILMFWY |
| 123.0 | 53.4 | 174.0 | 1111.1 | 1111.1 | 1111.1 | 0.4 | 0.3 | 0.4 | 0.4 | 0.4 | 0.4 | RDNQEHILMFWY |
| 124.3 | 54.9 | 175.9 | 1111.1 | 1111.1 | 1111.1 | 0.4 | 0.3 | 0.3 | 0.4 | 0.4 | 0.4 | RDNQEHILMFWY |
| 124.3 | 55.1 | 174.2 | 1111.1 | 1111.1 | 1111.1 | 0.5 | 0.5 | 2.0 | 0.4 | 0.4 | 0.4 | RDNQEHILMFWY |
| 124.8 | 62.4 | 175.0 | 1111.1 | 1111.1 | 1111.1 | 0.8 | 0.4 | 0.4 | 0.4 | 0.4 | 0.4 | T |
| 124.9 | 54.0 | 170.6 | 1111.1 | 1111.1 | 1111.1 | 0.4 | 0.3 | 1.0 | 0.4 | 0.4 | 0.4 | RDNQEHILMFWY |
| 125.4 | 54.0 | 174.9 | 1111.1 | 1111.1 | 1111.1 | 0.4 | 0.4 | 0.4 | 0.4 | 0.4 | 0.4 | RDNQEHILMFWY |
| 125.4 | 53.3 | 174.4 | 1111.1 | 1111.1 | 1111.1 | 0.4 | 0.4 | 0.4 | 0.4 | 0.4 | 0.4 | RDNQEHILMFWY |
| 126.2 | 52.7 | 173.9 | 1111.1 | 1111.1 | 1111.1 | 0.4 | 0.5 | 1.0 | 0.4 | 0.4 | 0.4 | RDNQEHILMFWY |
| 126.4 | 54.9 | 172.6 | 1111.1 | 1111.1 | 1111.1 | 0.4 | 0.3 | 0.3 | 0.4 | 0.4 | 0.4 | RDNQEHILMFWY |
| 126.4 | 55.3 | 174.0 | 1111.1 | 1111.1 | 1111.1 | 0.4 | 0.4 | 0.8 | 0.4 | 0.4 | 0.4 | RDNQEHILMFWY |
| 126.8 | 50.5 | 172.1 | 1111.1 | 1111.1 | 1111.1 | 0.6 | 0.4 | 0.4 | 0.4 | 0.4 | 0.4 | A |
| 127.1 | 52.7 | 172.0 | 1111.1 | 1111.1 | 1111.1 | 0.4 | 0.3 | 0.3 | 0.4 | 0.4 | 0.4 | RDNQEHILMFWY |
| 127.5 | 53.5 | 174.3 | 1111.1 | 1111.1 | 1111.1 | 0.4 | 0.4 | 0.4 | 0.4 | 0.4 | 0.4 | RDNQEHILMFWY |
| 128.8 | 64.5 | 171.3 | 1111.1 | 1111.1 | 50.3 | 0.4 | 0.4 | 0.4 | 0.4 | 0.4 | 0.4 | P |
| 129.4 | 50.6 | 172.4 | 1111.1 | 1111.1 | 1111.1 | 0.6 | 0.4 | 0.8 | 0.4 | 0.4 | 0.4 | A |
| 129.7 | 55.1 | 173.0 | 1111.1 | 1111.1 | 1111.1 | 0.4 | 0.3 | 0.3 | 0.4 | 0.4 | 0.4 | RDNQEHILMFWY |
| 129.9 | 62.6 | 171.8 | 1111.1 | 1111.1 | 1111.1 | 0.4 | 0.3 | 0.3 | 0.4 | 0.4 | 0.4 | P |
| 130.4 | 55.0 | 174.7 | 1111.1 | 1111.1 | 1111.1 | 0.4 | 0.3 | 0.3 | 0.4 | 0.4 | 0.4 | RDNQEHILMFWY |
| 133.1 | 55.5 | 173.8 | 1111.1 | 1111.1 | 1111.1 | 0.4 | 0.4 | 0.8 | 0.4 | 0.4 | 0.4 | RDNQEHILMFWY |
